# Modular engineering of embryonic-extraembryonic interactions generates advanced gastruloid morphotypes

**DOI:** 10.1101/2025.11.09.687163

**Authors:** Natalia P. Smirnova, Sergey V. Ponomartsev, Tharvesh M. Liyakat Ali, Max Lycke, Brian K. Chung, Jonas Øgaard, Espen Melum, Thomas Combriat, Jesse V. Veenvliet, Stefan Krauss Deiml

## Abstract

During mammalian embryogenesis, reciprocal interactions between the epiblast and extraembryonic endoderm are critical for germ layer specification and body plan development during gastrulation. Gastruloids recapitulate aspects of gastrulation in the absence of extraembryonic cues, resulting in a predominantly posteriorized and dorsalized phenotype with a limited lineage diversity. Here, we establish a modular co-aggregation (“aggregoid”) strategy that spatially couples embryonic and extraembryonic endoderm-like cells to reconstruct key interactions *in vitro*. This drives self-organized anterior-ventral patterning together with the emergence of node- and notochord-like structures, enriched endodermal populations, and increased mesodermal diversity, including cardiopharyngeal lineages and vascular endothelium. Our findings demonstrate that modular engineering of lineage interactions can direct self-organized patterning in stem-cell-based embryo models and provide a versatile framework for generating defined morphotypes.

## INTRODUCTION

Canonical mouse 3D gastruloids, a subclass of stem-cell-based embryo models (SCBEMs) recapitulate key features of embryonic development including axial organization, the emergence of the three germ layers (ectoderm, mesoderm and endoderm) and the initiation of organogenesis(Beccari et al., 2018; Brink and van Oudenaarden, 2021; Turner et al., 2017; Turner and Martinez Arias, 2024; van den Brink et al., 2014; Veenvliet et al., 2020; Veenvliet et al., 2021). This cascade of events, recapitulating the outcome of gastrulation, is triggered by transient and uniform exposure of mouse embryonic stem cell (mESC) aggregates to the WNT agonist CHIR99021 (CHIR)(Beccari et al., 2018; Brink and van Oudenaarden, 2021; Turner et al., 2017; Turner and Martinez Arias, 2024; van den Brink et al., 2014; Veenvliet et al., 2020; Veenvliet et al., 2021). However, *in vivo*, mammalian gastrulation is orchestrated by precise spatiotemporal deployment of multiple signaling pathways (e.g., WNT, BMP, NODAL, FGF, HH), with crucial contributions from the extraembryonic tissues that are absent in gastruloids(Bardot and Hadjantonakis, 2020; Haantjes et al., 2025; Morgani and Hadjantonakis, 2020). Consequentially, despite the striking ability for reproducible self-organization in the absence of instructive extra-embryonic cues, gastruloids exhibit limited lineage representation, including lack of anterior neural structures, and variable potential towards endodermal and anterior mesodermal fates(Beccari et al., 2018; Turner et al., 2017). This limitation can be explained by the uniform exposure to CHIR, which biases development towards a WNT3A/BRA late primitive streak (PS) program associated with caudal fates, and insufficient induction of the NODAL/EOMES-dependent early PS program required for specific mesendodermal and anterior identities(Arias et al., 2025; Dias et al., 2025; Wehmeyer et al., 2025). Moreover, the absence of critical signaling centers such as a mature notochord, results in a dorsalized organization of canonical gastruloids(van den Brink et al., 2014; Veenvliet et al., 2020).

Extraembryonic endoderm (ExEnd), including primitive endoderm (PrE) and its derivatives such as visceral endoderm (VE), anterior visceral endoderm (AVE) and parietal endoderm (ParE) play a multifaceted and dynamic role in supporting epiblast maturation and axis formation(Kumar et al., 2015), regulating PS progression(Tortelote et al., 2013; Yoon et al., 2015), and providing structural support, and transport of nutrients(Nowotschin and Hadjantonakis, 2020). A subpopulation of the VE, termed anterior visceral endoderm (AVE) is a critical signaling center that spatially restricts PS formation by expressing WNT antagonists like DKK1, and TGFβ/NODAL antagonists such as LEFTY1 and CERBERUS1 (CER1), thereby defining the anterior region of the embryo(Kumar et al., 2015). The posterior visceral endoderm (PVE), together with BMP4-expressing extra-embryonic ectoderm and the adjacent epiblast, induces WNT3, which initiates PS formation and gastrulation(Tortelote et al., 2013; Yoon et al., 2015).

Here, we present a versatile and modular co-aggregation strategy using transgene-free mESC-derived ExEnd-like cells to delineate the effects of the extraembryonic environment on lineage specification, patterning, and morphogenesis. We find that the presence of three alternative classes of ExEnd-like cells differentially modulates the balance of NODAL and WNT signaling promoting PS regionalization, lineage diversity, and increased representation of endoderm, axial mesoderm, and cardiopharyngeal lineages, while enhancing ventral and anterior features. Overall, our aggregation approach provides versatile platform for engineering SCBEM composition and developmental potential.

## RESULTS

### Derivation of ExEnd-like cells

To generate gastruloid-based mouse SCBEMs containing an ExEnd compartment, we adapted published protocols for the derivation of cell types resembling pre- and post-implantation ExEnd-like cells from mESCs without induced transgene overexpression. Previous studies(Anderson et al., 2017; Perera et al., 2022) showed that culturing naïve mESCs (2i cells) in RACL (RPMI supplemented with B27 without insulin, Activin A, CHIR, Lif (Methods)) medium induces differentiation into a PrE-like state, which can be further converted into a VE-like identity in NACLB (N2B27, Activin A, CHIR, Lif, BMP4) medium (Figure 1A, Methods). Next, based on findings that WNT inhibition facilitates an AVE-like transcriptional profile(Schumacher et al., 2024), we replaced CHIR in NACL with the WNT pathway inhibitor XAV939 to generate XAL (N2B27, XAV939, ActivinA, Lif) medium (Figure 1A, Methods). RACL, NACLB and XAL identities were assessed by marker gene analysis (Supplementary 1A) and comparison to the integrated datasets from published mouse embryonic atlases(Argelaguet et al., 2019; Cheng et al., 2019; Liu et al., 2022; Mohammed et al., 2017; Thowfeequ et al., 2024) (Supplementary 1B).

**Figure 1.**
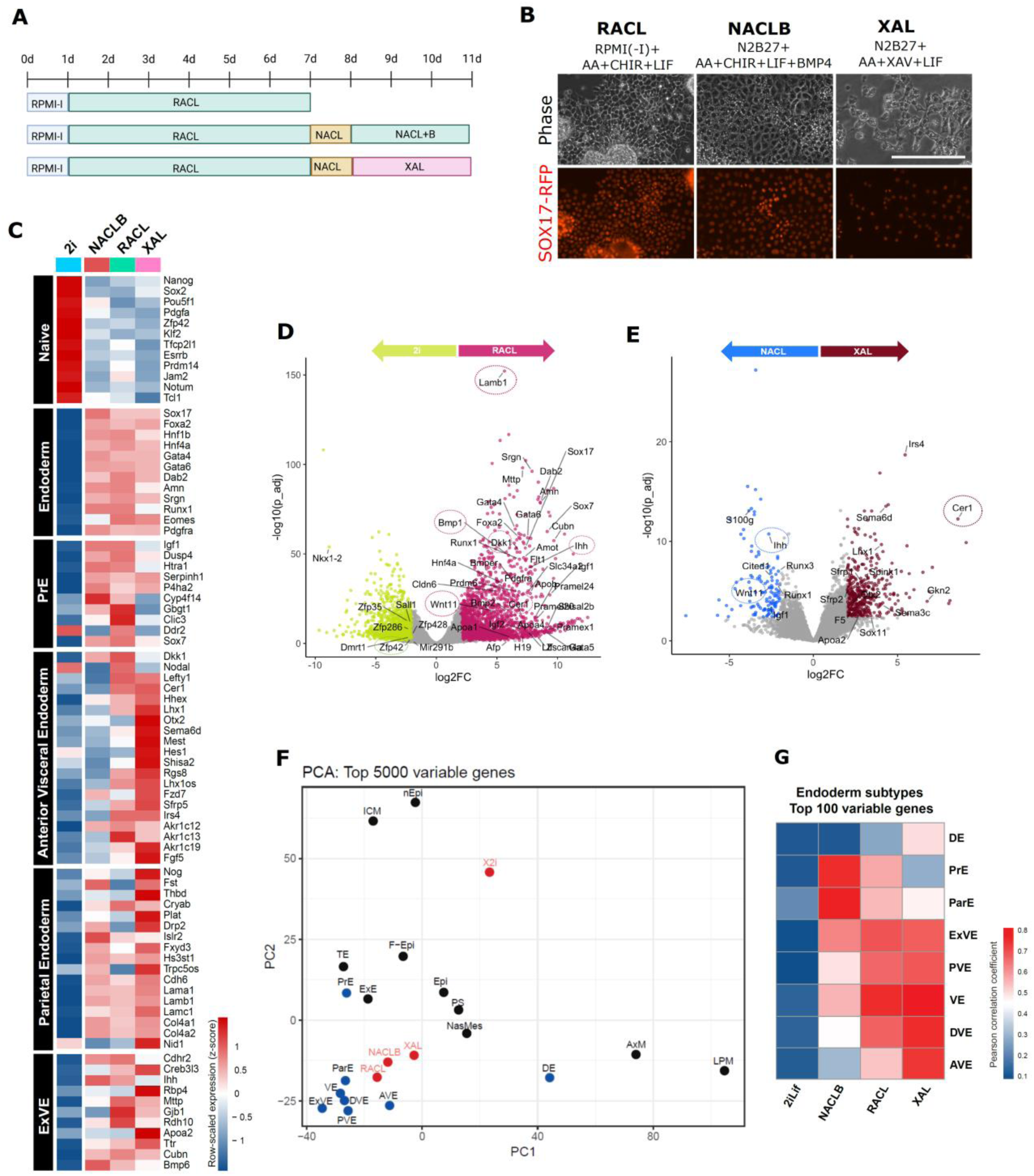
Derivation of ExEnd-like cells. (A) Outline of the derivation of ExEnd-like cell types from mESC grown in 2i/N2B27. (B) Representative phase contrast and *Sox17*-RFP reporter expression images of cells in RACL, NACLB and XAL conditions at the end point of the differentiation protocol. Scale bar: 100 µm. (C) Heatmap of naïve pluripotency and extraembryonic endoderm marker gene expression across 2i/N2B27, RACL, NACLB and XAL conditions. Data are row-wise z-score normalized (n=3 biological replicates). The marker list was derived from a differential expression analysis of integrated published datasets(Argelaguet et al., 2019; Cheng et al., 2019; Liu et al., 2022; Mohammed et al., 2017; Thowfeequ et al., 2024) (Supplementary 1A). (D and E) Volcano plot of differential gene expression comparing 2i/N2B27 and RACL cells (D) and NACLB and XAL cells (E). (F) PCA of bulk RNA-seq profiles from 2i/N2B27, RACL, NACLB and XAL cells (n=3) integrated with pseudo-bulked reference datasets from mouse embryonic cell types (Supplementary 1B) based on the top 5000 variable genes. (G) Heatmap showing correlation scores between bulk RNA-seq profiles of 2i/N2B27, RACL, NACLB, and XAL cells (n = 3) and selected cell types from integrated reference datasets.

Conversion of 2i cells towards ExEnd-like states was accompanied by morphological changes along with SOX17, GATA4 and PDGFRα expression (Figure 1B, Supplementary 1C). Bulk RNA-sequencing and quantitative RT-PCR (qPCR) analyses demonstrated a loss of a naïve pluripotency transcriptional signature of 2i cells and acquisition of a *Gata6, Gata4, Hnf4a, Pdgfra, Lama1* and *Ttr-*positive pan-ExEnd expression profile in all three conditions (Figure 1C-D, Supplementary 1D). RACL cells displayed enrichment for markers associated with both PrE (*Igf1, Sox7* and *Dusp4*) and VE (*Dab2, Cubn, Afp, Apoa1*) (Figure 1C-D, Supplementary 1E) together with a distinct cluster of transcripts linked to pluripotency, germline and early embryogenesis(Srinivasan et al., 2020) including *Zscan4, and Pramel16* (Supplementary 1E). Upon further differentiation of RACL cells in NACLB and XAL conditions, cells developed lipid droplets, consistent with features of ExEnd maturation(Artus et al., 2012) (Figure 1B). Notably, we did not observe a substantial downregulation of markers that are associated with a PrE-to-VE transition such as *Sox7(Artus et al., 2012; Weberling et al., 2025)* or *Sox17* in NACLB cells (Figure 1C). XAL cells displayed a OTX2, LEFTY1, and EOMES-positive profile, consistent with an AVE-like identity (Supplementary 1F, G). Bulk RNA-Seq and qPCR showed an up-regulation of AVE-associated genes, including *Hhex, Lhx1, and Otx2* and secreted antagonists such as *Lefty1, Dkk1, and Cer1* (Figure 1C, Supplementary 1H, I). Among XAL-specific transcripts were also genes linked to cell migration such as *Sema6* and *Sox11* which have been associated with AVE(Thowfeequ et al., 2024)(Figure 1E). The expression of *Dkk1* was significantly increased in both RACL and NACLB cells, but not in XAL (Figure 1C, Supplementary 1H). *Cer1* and *Lefty1* were only expressed in XAL and RACL cells (Figure 1C, Supplementary 1H). RACL and NACLB cells also expressed genes encoding the non-canonical WNT ligand *Wnt11* and the HH pathway ligand *Ihh*, together with extracellular matrix components such as *Lama1 and Lamb1* (Figure 1D-E, Supplementary 1I).

A principal component analysis (PCA) using the top 5000 variable genes, benchmarked against reference mouse E3.5-7.5 embryonic datasets(Argelaguet et al., 2019; Cheng et al., 2019; Liu et al., 2022; Mohammed et al., 2017; Thowfeequ et al., 2024), grouped all three conditions with ExEnd cell types, but distinct from definitive endoderm (DE) (Figure 1F). Correlation analysis based on the top 100 variable genes indicated that RACL cells most closely resembled VE, while NACLB cells correlated with ParE and PrE, while XAL showed highest similarity to distal visceral endoderm (DVE) and AVE (Figure 1G). Notably, RACL cells also exhibited partial similarity to AVE/DVE populations and had a high correlation score with DVE (Figure 1G). Overall, the obtained ExEnd-like populations displayed a molecular signature consistent with an ExEnd-like identity albeit with limitations characteristic for *in vitro* ExEnd derivation protocols(Anderson et al., 2017; Julio et al., 2011; Kunath et al., 2005; Niakan et al., 2013; Ohinata et al., 2022; Perera and Brickman, 2023; Shimosato et al., 2007; Wamaitha et al., 2015; Zhong et al., 2018). Importantly, a diverse repertoire of signaling molecules in the ExEnd-like cells suggested their potential in axis establishment and lineage regulation(Hoshino et al., 2015; Kretzschmar et al., 2025).

### ExEnd co-aggregation promotes early pre-patterning and symmetry breaking

To investigate the influence of ExEnd-like cells on early patterning of mouse SCBEMs, we co-aggregated mESCs with either of the three pre-differentiated ExEnd-like cell pools in a proportion of 300:100 cells respectively. A pulse of CHIR (3 µM) was applied between 48 and 72h, as in the canonical gastruloid protocol(Beccari et al., 2018; van den Brink et al., 2014) (Figure 2A). The resulting structures were termed RACL-, NACLB-, and XAL-aggregoids, depending on the specific cell types that were used for the co-aggregation, and collectively referred to as ExEnd-aggregoids. Structures formed from mESCs cells alone were referred to as control gastruloids.

**Figure 2.**
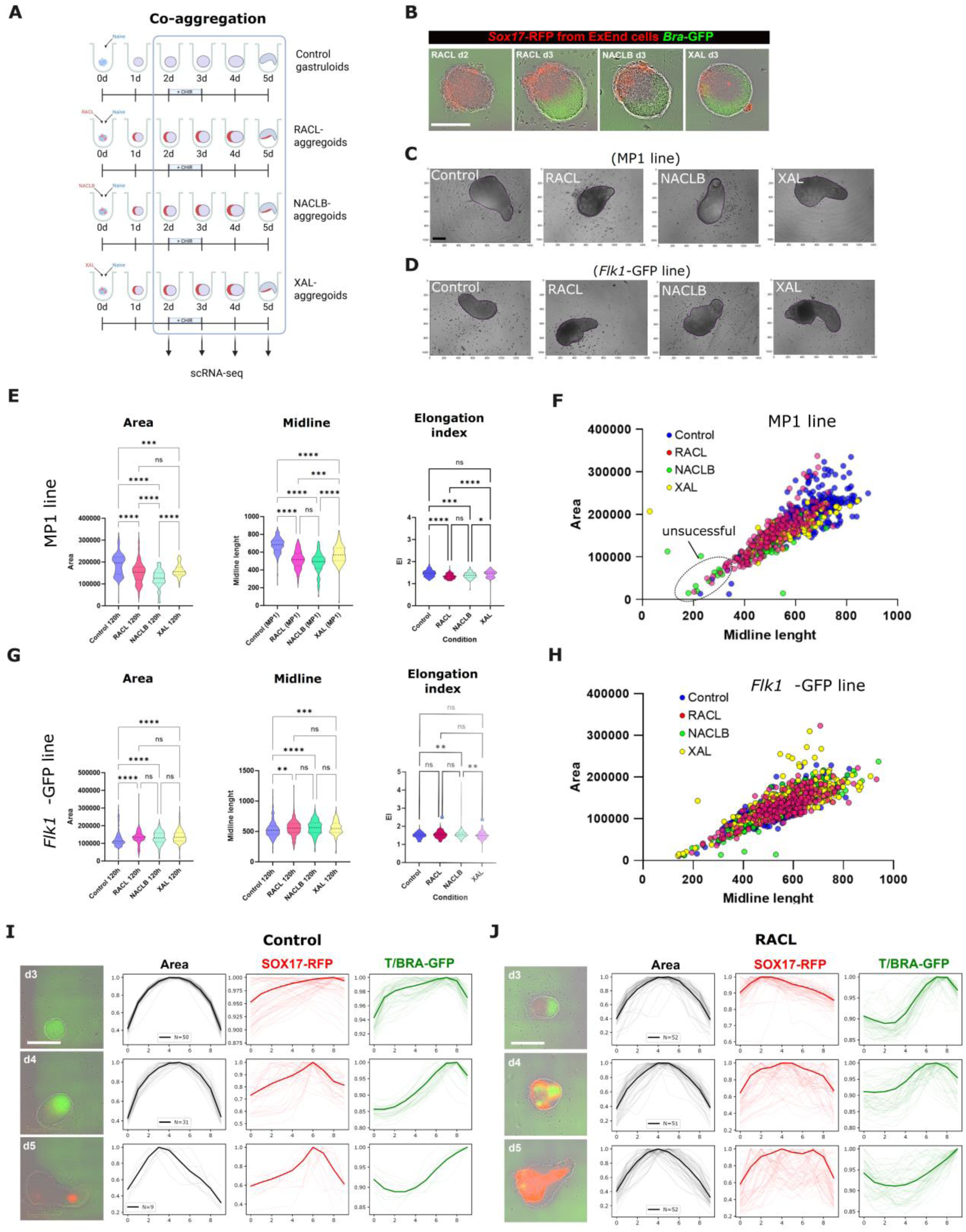
ExEnd co-aggregation promotes early pre-patterning and symmetry breaking. (A) Experimental outline and sampled time points of the co-aggregation protocols. (B) Representative phase contrast images with overlayered green (*Bra*-GFP reporter) and red (*Sox17*-RFP from added ExEnd-like cells) fluorescent signals of 72h ExEnd-aggregoids demonstrating the “cap” of *Sox17*-RFP (derived from ExEnd-like cells) and the polarized *Bra*-GFP reporter after CHIR pulse. Scale bar: 200 µm. (C-D) Representative brightfield images of 120h control gastruloids, RACL-, NACLB- and XAL-aggregoids made of MP1 (C) and *Flk1*-GFP (D) with automatically outlined measured area. Scale bar: 200 µm. (E, G) Violin plots demonstrating measurements of area, midline length and Elongation index of control gastruloids and ExEnd-aggregoids made of MP1 (n(control)=311, n(RACL)=259, n(NACLB)=82, n(XAL)=101)(F) and *Flk1*-GFP (n(control)=330, n(RACL)=250, n(NACLB)=378, n(XAL)=287)(H) lines at 120h. Kruskal-Wallis test. (F, H) Scatter plots showing correlation of midline length and area of gastruloids and aggregoids made of MP1 line, n=687 (E) and Flk1-GFP line, n=1361 (G). (I-J) Phase contrast images with overlayered green (*Bra*-GFP reporter) and red (*Sox17*-RFP) fluorescent signals with optical segment segregated measurements of area, GFP, and RFP intensity (individual values and mean). Control gastruloids (I) and RACL-aggregoids (J). Scale bar: 400 µm.

Upon co-aggregation, successful symmetry breaking and elongation was observed in 87% of RACL- (n=1859, 19 experimental replicates), 67% of NACLB- (n=616, 10 experimental replicates) and 98% of XAL- (n=923, 10 experimental replicates) aggregoids (Supplementary 2A-C). “Unsuccessful” structures that were fully enveloped by ExEnd-like cells, failed to grow and elongate (Supplementary 2D). Nevertheless, by 120h these structures exhibited a SOX17- and BRA-positive phenotype, indicating that full coverage with ExEnd-like cells can suppress growth and elongation while still permitting lineage specification.

“Successful” aggregoids were partially covered by ExEnd-like cells which localized to the surface at 48-72h (Figure 2B) and resembled a general gastruloid morphology by 120h (Figure 2C-D). Morphometric measurements showed that, depending on the cell line used, 120h RACL- and NACLB-aggregoids exhibited either slightly reduced area, midline length and elongation indexes when using *Bra*-GFP: *Sox17*-RFP double reporter MP1(Pour et al., 2022) line (Figure 2E) or comparable parameters in *Flk1*-GFP line (Figure 2G) relative to control gastruloids. The elongation index of XAL-aggregoids did not significantly differ from controls in both cell lines (Figure 2E-G). The projected area and midline length were positively correlated and broadly overlapping between conditions for both cell lines, indicating conserved scaling between growth and axial extension (Figure 2F-H). However, the scatter plot also revealed a subset of low-area, low-midline length outliers, predominantly seen in NACLB- and RACL-aggregoids (Figure 2F), whereas long-midline outliers with increased area and midline length were mainly observed in the control gastruloid group (MP1 line) (Figure 2F) or XAL-aggregoids (*Flk1*-GFP line) (Figure 2H). Overall, the capacity for axial elongation was retained in all experimental conditions.

To visualize symmetry breaking dynamics, we used a cell line with *Bra*-GFP (marking PS and early mesoderm) and *Sox17*-RFP (marking endoderm) fluorescent reporters, further referred to as *Bra*-GFP: *Sox17*-RFP (MP1 line) together with time-lapse microscopy and an algorithm for fluorescence intensity quantification along the longitudinal axis of the structure (Supplementary 2E-F, Methods).

Following CHIR stimulation, *T/Bra*-GFP expression in control gastruloids displayed a radially symmetric pattern at 72h (Figure 2I, Movie 01). In contrast, ExEnd-aggregoids exhibited polarized *T/Bra*-GFP expression from the onset, expanding from the pole opposite the ExEnd cap (Figure 2G, I, Movie 02). Because ExEnd-like cell types expressed the WNT inhibitor *Dkk1* (Supplementary 1H), we hypothesize that this cap could locally attenuate CHIR-induced WNT signaling and downstream T/BRA activation. Quantification of GFP intensity along the anterior-posterior axis confirmed uniform *T/Bra*-GFP distribution in controls (Figure 2I) and a localized peak in RACL-aggregoids (Figure 2J) at 72h. At 96h, the *T/Bra*-GFP domain in controls gradually localized to the posterior pole forming a sharp intensity peak across optical segments 6-8 (Figure 2I), while in RACL-aggregoids it extended anteriorly, generating a broader and smoother distribution profile at 96-120h (Figure 2J), often with the emergence of a secondary peak at the anterior pole (Supplementary 2E-F).

Together, these data demonstrate that ExEnd co-aggregation reshapes symmetry breaking and the dynamics of axis formation in mSCBEMs while remaining compatible with a gastruloid-like elongation.

### ExEnd co-aggregation alters cell composition in mSCBEM

To assess the effects of ExEnd co-aggregation on mSCBEM cell composition, we performed single-cell RNA-sequencing (scRNA-seq) across all four conditions (control gastruloids, RACL-, NACLB-, XAL-aggregoids) at 48h (pre-CHIR), 72h (post-CHIR), 96h and 120h (maximum elongation). Cell types were annotated by label transfer from a reference mouse embryo E6.5-E9.5 atlas(Imaz-Rosshandler et al., 2024; Pijuan-Sala et al., 2019) (Supplementary 3A-B), followed by filtering and manual merging based on prediction scores, cluster size and gene expression (Figure 3A-D, Supplementary 3C-F).

**Figure 3.**
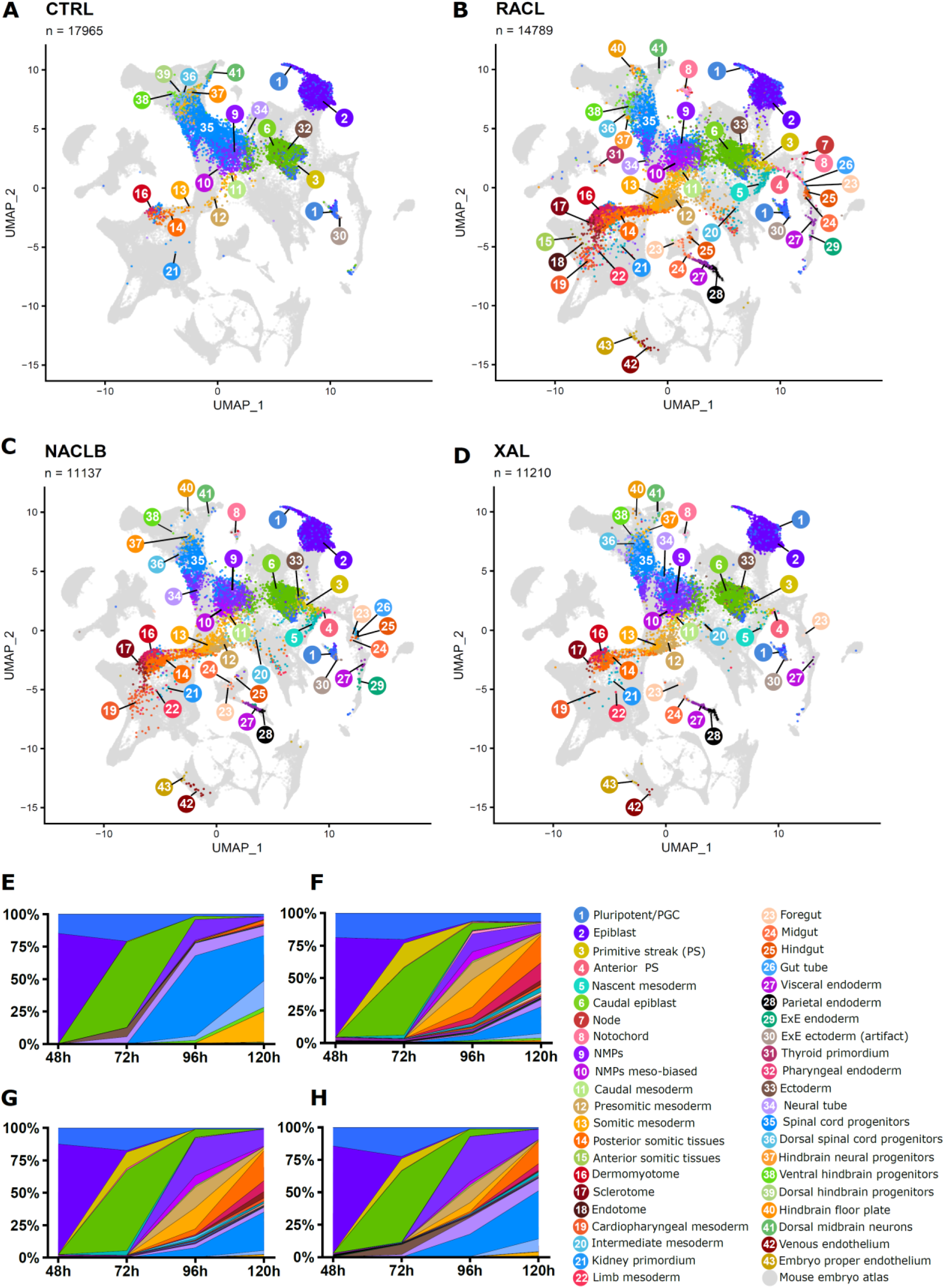
ExEnd co-aggregation alters cell composition in mSCBEM. (A-D) UMAP visualization of control gastruloids and RACL-, NACLB- and XAL-aggregoids, colored by cluster identities. The control gastruloid and ExEnd-aggregoid datasets are integrated with a mouse E6.5-9.5 dataset(Imaz-Rosshandler et al., 2024; Pijuan-Sala et al., 2019) (Supplementary 3A-B). The color code is shown below. (E-H) Stacked overflow plots demonstrating cell composition of control gastruloids and ExEnd-aggregoids color-coded by cluster’s identity.

Across the 48-120h time course, all conditions showed progression from pluripotent states before CHIR pulse towards an exit of pluripotency and the emergence of lineage-committed populations after CHIR treatment (Figure 3E-H). By 96h, bipotential neuromesodermal progenitors (NMPs) co-expressing *Sox2* and *Bra*(Binagui-Casas et al., 2025; Koch et al., 2017) (Figure 3E-H, Supplementary 3K) emerged followed by neural and mesodermal lineage diversification. Control gastruloids displayed a pronounced neuroectodermal bias by 120h (Figure 3E). RACL- and NACLB-aggregoids showed higher proportion of *T/Bra*-high (Figure 3F-G) mesoderm-biased(Gouti et al., 2017) NMPs (37% and 23%) compared to control and XAL conditions (5% and 8%) (Figure 3E, H), which was associated with an increased mesodermal output and reduced neuroectodermal bias at later stages. By 120h, the composition of control gastruloids was represented by paraxial mesoderm (presomitic mesoderm, somitic mesoderm, posterior somitic tissues, and dermomyotome) and neuroectodermal clusters (neural tube, dorsal midbrain progenitors, dorsal spinal cord progenitors, hindbrain neural progenitors, dorsal and ventral hindbrain progenitors) with the sufficient neural bias (Figure 3E). ExEnd-aggregoids demonstarated higher lineage diversity, additionally containing cell types that were absent or underrepresented in controls, including ExEnd populations (ExEnd, VE, ParE), cardiopharyngeal mesoderm and endothelial clusters (embryo proper endothelium, venous endothelium), notochord, DE (gut tube, foregut, midgut, hindgut), and also ventral-associated cell types such as hindbrain floor plate, sclerotome and endotome (Figure 3F-H). Those cell types were best represented in RACL-aggegoids, showed a comparable enrichment in NACLB-aggregoids, but were less present in XAL-aggregoids (Figure 3B-D, F-H).

To investigate the origin of cell diversity and biases in control gastruloids and aggregoids, we performed trajectory inference using Monocle3(Trapnell et al., 2014). At 72h, the developmental trajectory in control gastruloids exhibited two principal branches: one progressing through caudal epiblast (CE) to NMPs, generating spinal cord progenitors and paraxial mesoderm (PM), and a second one enriched for pluripotent cell types, ectoderm and neural tube (NT) populations contributing to neuroectoderm (Supplementary 3C, G). In contrast, RACL-aggregoids displayed a dual origin of mesoderm. One trajectory branch followed the NMP route like in controls, while a second emerged from pluripotent populations, passed through PS and nascent mesoderm (NM), and contributed to paraxial, lateral plate and intermediate mesoderm lineages (Supplementary 3D, H). A similar pattern was observed in NACLB (Supplementary 3 E, I) but less so in XAL (Supplementary 3F, J). This is consistent with previous reports describing multiple mesodermal origins arising from both PS- and NMP-associated trajectories(Guibentif et al., 2021) as well as the emergence of cardiac mesoderm from the early PS(Costello et al., 2011).

Overall, co-aggregation with ExEnd-like cells was associated with an increase in cell type diversity and a shifted lineage composition.

### ExEnd co-aggregation promotes early PS regionalization and lineage diversification

Co-aggregation with ExEnd-like cells promoted DE and anterior mesoderm (AM) - lineages associated with the early PS and NODAL/EOMES-dependent program typically underrepresented in control gastruloids which mostly employ WNT3a/BRA regulatory module(Arnold et al., 2008; Dias et al., 2025; Probst et al., 2021; Schule et al., 2023; Wehmeyer et al., 2025). To characterize of condition-specific differences in trajectory deployments, we first focused on the time points before (48h) and after (72h) CHIR pulse.

At 48h, both control gastruloids and ExEnd-aggregoids were predominantly composed of cells expressing *Otx2, Fgf5, Utf1, Rasgrp2,* and *Dppa5a*, consistent with a pre-gastrulation epiblast state (Figure 4A-B, D, together with a population annotated as “PGC/Pluripotent” corresponding to a range of pluripotency states (Figure 4C-D). At this stage, very few cells expressed markers that indicate pluripotency exit (Figure 4D), and aside from the presence of ExEnd cells in aggregoids, no substantial differences in cell type proportions, marker expression or WNT/NODAL module scores were detected (Figure 4A-B, Supplementary 4A-B), suggesting that major condition-specific differences emerge after 48 h. However, an intercellular communication analysis by CellChat(Jin et al., 2025) revealed ligand-receptor pairs associated with laminin, collagen, and BMP signaling between ExEnd and epiblast, which were absent in controls and XAL-aggregoids (Supplementary 4C).

**Figure 4.**
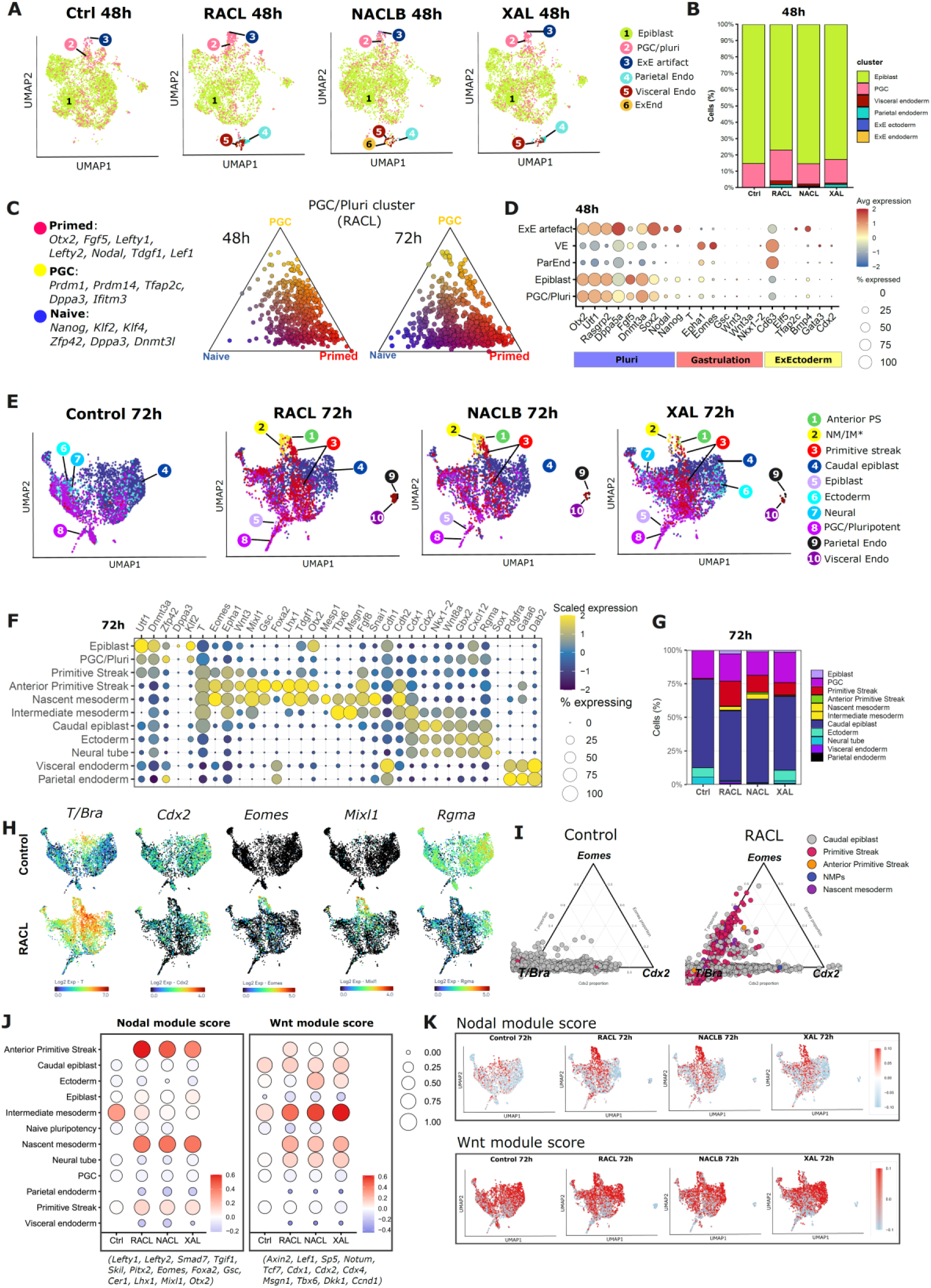
ExEnd co-aggregation promotes early PS regionalization and lineage diversification. (A) UMAP showing annotated cell clusters in control gastruloids and RACL-, NACLB-, and XAL-aggregoids at 48 h, based on label transfer from a mouse embryo reference atlas(Imaz-Rosshandler et al., 2024; Pijuan-Sala et al., 2019). Clusters with fewer than 10 cells were excluded. (B) Stacked bar plots showing proportions of annotated cell types at 48 h. Clusters with fewer than 10 cells were excluded. (C) Ternary plots showing naïve, primed, and PGC-like scores within the automatically annotated “PGC/Pluri” cluster at 48 h and 72 h. (D) Dot plot showing expression of markers associated with pluripotency, exit from pluripotency/gastrulation, and extraembryonic lineages at 48 h. The cluster initially annotated as “ExE ectoderm” shows low expression of canonical markers and likely represents misannotated undifferentiated cells. (E) UMAP showing annotated cell clusters at 72 h across all conditions. Clusters with fewer than 10 cells were excluded. (F) Dot plots showing marker gene expression in annotated cell types at 72 h. (G) Stacked bar plots showing proportions of annotated cell types at 72 h. Clusters with fewer than 10 cells were excluded. (H) UMAP projections of selected gene expression in control gastruloids and RACL-aggregoids at 72 h. (I) Ternary plots showing relative expression of *T/Bra*, *Cdx2*, and *Eomes* in control gastruloids and RACL-aggregoids at 72 h, colored by annotated cell types. (J) Dot plots showing NODAL and WNT module scores across annotated clusters in all conditions. Module scores were calculated based on predefined gene sets (see Methods). (K) UMAP feature plots showing NODAL and WNT module scores at 72 h across all conditions.

By 72h, at the scRNA-seq level, only 54% of cells in control gastruloids expressed *T/Bra*, compared to 89%, 91%, and 84% in RACL-, NACLB-, and XAL-aggregoids, respectively. However, this pattern was consistent with the start of a posterior restriction of the *Bra-*GFP-positive domain in controls and its anterior expansion in ExEnd-aggregoids at 84-96h. At this stage, *T/Bra* and *Cdx2*-positive CE (Figure 4F, H, I) dominated in cell composition of both control gastruloids and ExEnd-aggregoids (Figure 4E, G). However, ExEnd-aggregoids contained *T/Bra* and *Eomes* double-positive cell types (Figure 4F, H, I) absent or underrepresented in controls, annotated as PS, Anterior PS (APS), and NM (Figure 4E, G). NM and APS shared *Eomes, Mixl1, Lhx1, Tdgf1, Otx2* and *Gsc*-positive signature (Figure 4F, H), but could be distinguished by *Foxa2* expression in APS and *Mesp1* in NM (Figure 4F), consistent with published data(Costello et al., 2011; Masamsetti et al., 2025; Osteil et al., 2025; Probst et al., 2021). Moreover, control gastruloids and XAL-aggregoids contained *Rgma*-positive populations (Figure 4H) annotated as ectoderm and NT, suggesting an early neural bias (Figure 4G). To further characterize pathways activity, we computed NODAL and WNT module scores and projected them onto the UMAP space. Overall, RACL- and NACLB-aggregoids exhibited higher NODAL module scores, particularly within APS, PS, and NM populations. (Figure 4J-K, Supplementary 4E-F). WNT module activity was also observed in ExEnd-aggregoids, particularly in emerging mesodermal populations. (Figure 4J-K, Supplementary 4E). Analysis of module scores across multiple pathways also revealed increased FGF module activity in all ExEnd-aggregoids, as well as upregulated Notch module in RACL- and NACLB-aggregoids (Supplementary 4E-F).

These observations suggest that ExEnd co-aggregation extends the window of PS-like states, enabling the coexistence of early (*Eomes-*associated) and late (*Cdx2*-associated) transcriptional programs(Amin et al., 2016; Arnold et al., 2008) within the same structure (Figure 4I) leading to an activation of NODAL-associated module while preserving robust axial extension.

CellChat analysis at 72h showed that control gastruloids were dominated by BMP and WNT signaling, whereas ExEnd-aggregoids also exhibited enrichment of extra-cellular matrix (ECM) (with ExEnd as a major sender), FGF and VEGF (from PS, NM and intermediate mesoderm (IM)) and TGFb/NODAL (primarily from NM) signaling (Supplementary 4D)

Altogether, these results suggest that in aggregoids, ExEnd-like derived cell populations reshape the signaling landscape following CHIR stimulation, contributing to early regionalization and diversification of cell states.

### ExEnd co-aggregation drives axial patterning

Given that scRNAseq revealed additional *T/Bra*-positive populations such as PS, APS and NM in ExEnd-aggregoids but not in control gastruloids, we focused on T/BRA expression domains as hallmarks of early axial patterning.

At 72h, upon the CHIR pulse, T/BRA domain displayed radial symmetry with no or few EOMES-positive cells distributed in central part of the structure (Figure 5A). In contrast, ExEnd-aggregoids exhibited a T/BRA domain at the future posterior pole, opposite the DKK1-positive ExEnd cap and the anterior SOX2-positive domain (Figure 5B-C). At this stage, EOMES co-localized with T/BRA in ExEnd-aggregoids, resembling an early PS signature(Schule et al., 2023) (Figure 5B-C). Together, these data indicated that co-aggregation with ExEnd-like cells was associated with earlier symmetry breaking and spatial patterning, consistent with localized modulation of WNT signaling.

**Figure 5.**
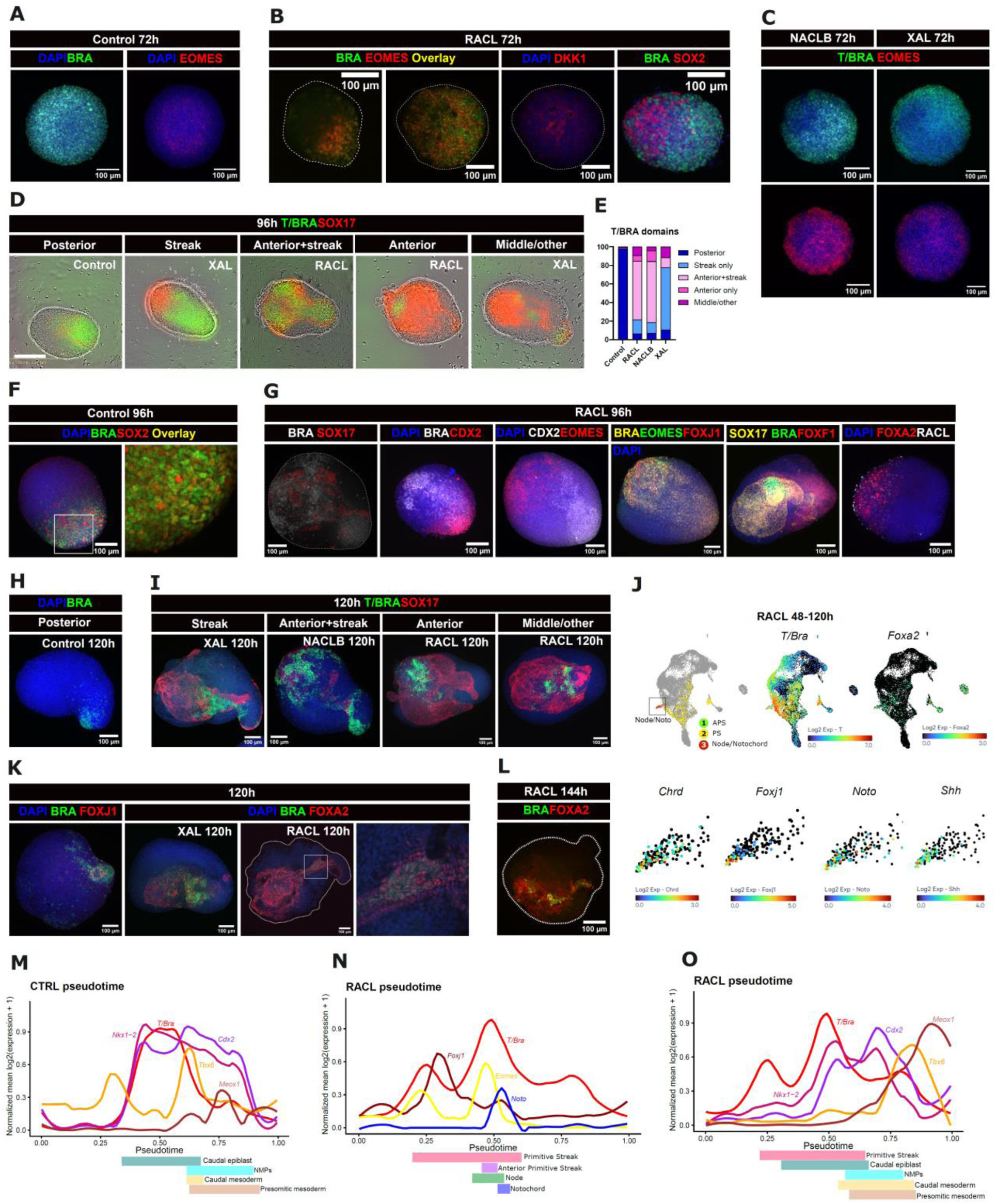
ExEnd co-aggregation drives axial patterning. (A) Representative confocal immunofluorescent images of T/BRA and EOMES expression in control gastruloid at 72h. (B) Representative confocal immunofluorescent images in RACL-aggregoids demonstrating localization of T/BRA, EOMES, DKK1, SOX2 at 72h. (C) Representative confocal immunofluorescent images demonstrating localization of T/BRA, EOMES in NACLB-and XAL-aggregoids at 72h. (D) Phase contrast images with overlayed green (*Bra*-GFP) and red (*Sox1*7-RFP) fluorescent signals demonstrating variability of T/BRA expression on gastruloids and ExEnd-aggregoids at 96h. Scale bar: 200 µm. (E) Bar plots demonstrating proportions of different morphology of T/BRA domains (n=1046) in control, RACL, NACLB- and XAL-conditions. (F) Representative confocal immunofluorescent image with magnified region showing SOX2 and T/BRA double-positive cells (NMPs) at 96h in control gastruloids. (G) Representative confocal immunofluorescent images of PS/APS/Axial mesoderm and early regionalization markers at 96h in RACL-aggregoids. (H) Representative confocal immunofluorescent image of a control gastruloid at 72h with a single T/BRA domain at posterior tip of the structure. (I) Representative confocal fluorescent images demonstrating the diversity and types of T/BRA domains in ExEnd-aggregoids (anterior, middle, midline/” streak”) at 120h. (J) UMAP projection of PS, APS and Node/Notochord clusters and axial mesoderm markers for RACL-aggregoids. (K) Representative confocal immunofluorescent images showing T/BRA (*Bra*-GFP reporter) and FOXJ1 or FOXA2 expression (marking axial mesoderm) at 120h. (L) Representative confocal immunofluorescent image showing T/BRA and FOXA2 expression (marking axial mesoderm) at 144h. (M-O) Smoothed expression dynamics of selected developmental marker genes plotted along normalized pseudotime trajectories in control (M) and RACL (N, O) conditions. Curves represent scaled expression intensity of lineage-associated genes across pseudotime, Colored bars below each plot indicate selected annotated cell states along the trajectories.

By 96h, control gastruloids showed progressive polarization and restriction of the T/BRA expression to the posterior tip, co-localizing with SOX2 consistent with an NMP-like identity(Wymeersch et al., 2016) (Figure 5D, F). At the same timepoint, ExEnd-aggregoids showed an anterior extension of the T/BRA domain, forming distinct midline/”streak” and anterior domains (Figure 5D) that persisted until 120h (Figure 5I) and were almost not observed in controls. Quantitively, 63% and 66% of RACL- and NACLB-aggregoids, respectively exhibited both anterior and “streak” domains, whereas the majority of XAL-aggregoids (67%) displayed only a “streak” T/BRA domain (Figure 5E). That was consistent with the comparatively lower representation of the APS cluster in XAL-aggregoids relative to RACL- and NACLB-aggregoids (Figure 4 E-G).

Immunostaining revealed regionalization of the T/BRA domain in RACL-aggregoids at 96h, with CDX2-positive posterior and EOMES-positive anterior compartments (Figure 5G). This region was also positive for FOXA2 supporting its APS identity (Figure 5G). The anterior region also contained a population of FOXJ1-positive cells, consistent with a node-associated identity (Figure 5G, J), which was later detected at more posterior positions (Figure 5K).

By 120h, the T/BRA signal in control gastruloids was localized at the posterior tip or diminished (Figure 5H). In contrast, ExEnd-aggregoids at 120-144h contained an elongated domain of T/BRA and FOXA2 double-positive cells consistent with a notochord-like identity (Figure 5I, K-L). In support, scRNA-seq analysis identified clusters expressing *Shh, Foxa2, Chrd*, and *Noto*, annotated as node/notochord-like populations (Figure 5J).

Using Monocle3, we reconstructed gene expression dynamics along a pseudotime axis to examine temporal relationships between T/Bra and axial patterning genes. In control gastruloids, *T/Bra* expression exhibited a single prominent peak at mid-pseudotime (∼0.5), coinciding with *Cdx2* and *Nkx1-2* expression and corresponding to CE states, followed by decline upon NMP emergence due to significant neural bias and poorly represented mesoderm (Figure 5M). RACL-aggregoids exhibited broader and temporally extended *T/Bra* expression, with multiple waves across the pseudotime (Figure 5N-O). First two waves were accompanied by a biphasic *Eomes* expression: an early peak consistent with PS-like activity and a later peak aligning with APS features and the onset of *Noto* expression, indicative of notochord specification (Figure 5N). A later smaller wave followed *Nkx1-2* and *Cdx2* peaks and coincided with *Tbx6* upregulation, consistent with caudal mesoderm differentiation (Figure 5O).

Together, these results suggest that co-aggregation with ExEnd-like cells extends and reshapes *T/Bra* dynamics and promotes the emergence of axial structures associated with an anterior-posterior organization.

### ExEnd co-aggregation promotes and organizes DE formation

In the mouse embryo, endoderm arises from two distinct sources: ExEnd, specified during pre-implantation, and DE which emerges during gastrulation from the APS(Nowotschin and Hadjantonakis, 2020), ingresses through the streak, displaces the VE, and gives rise to the gut epithelium and its derivatives(Ferrer-Vaquer et al., 2010).

*In vitro*, gastruloids show limited and unstable capacity to generate DE, showing substantial variability depending on the cell line and culture conditions(Arias et al., 2022; Blotenburg et al., 2025; Brink and van Oudenaarden, 2021; Farag et al., 2024; López-Anguita et al., 2022; Turner et al., 2017; Underhill and Toettcher, 2023). Control gastruloids derived from the *Flk1*-GFP line showed a pronounced deficiency of endoderm formation: at 120 h, SOX17-positive cells were sparse and failed to organize into continuous epithelial structures (Figure 6A), and endodermal clusters were not detected by scRNA-seq (Figure 3A). The MP1 cell line exhibited higher ability to produce SOX17-positive cells in controls but typically displayed a dispersed gut morphology(Farag et al., 2024) (Figure 6A).

**Figure 6.**
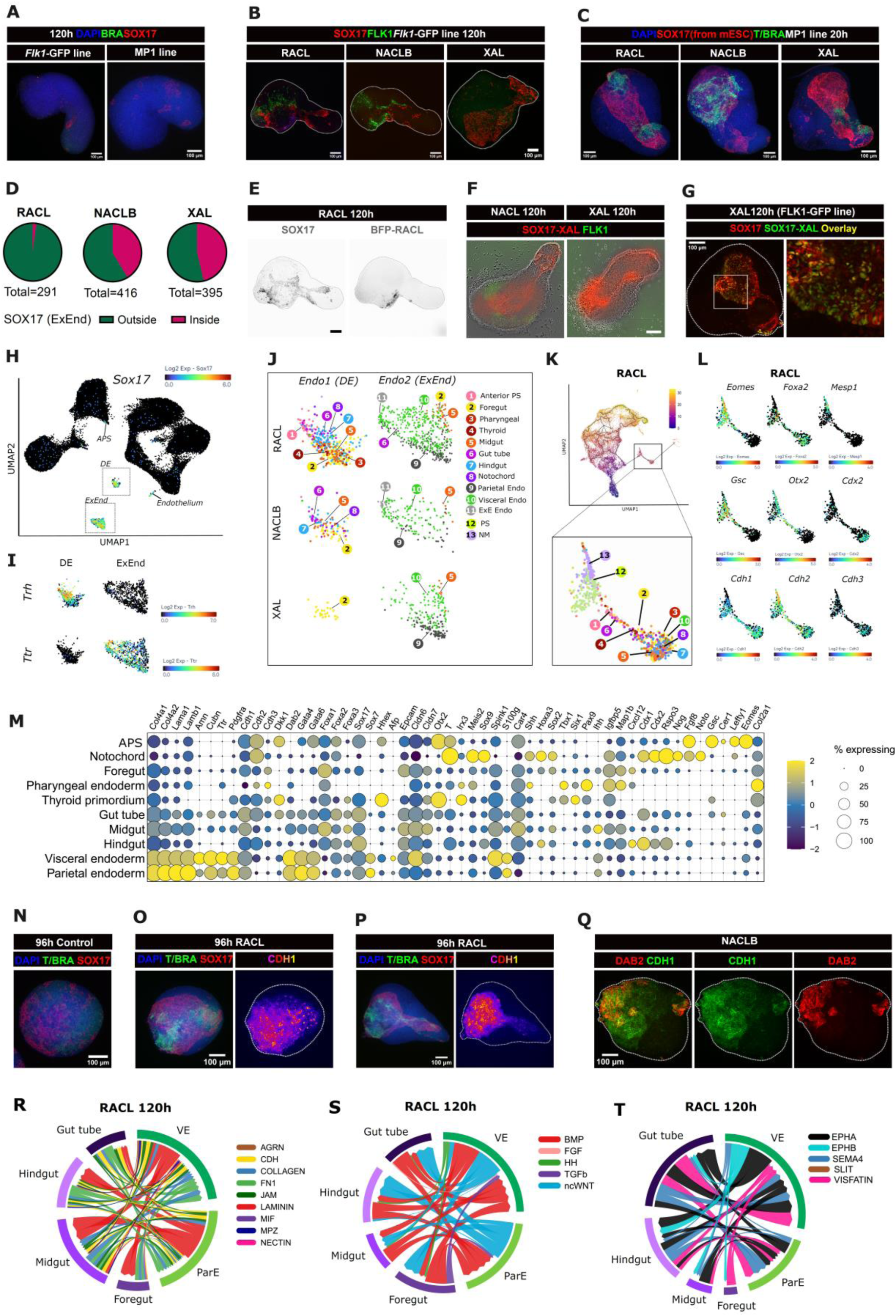
ExEnd co-aggregation promotes and organizes definitive endoderm formation. (A) Representative confocal immunofluorescent image of a 120h control gastruloids derived from the FLK1-GFP and MP1 lines, - and showing limited presence of SOX17 cells and lack of tube-like structure. (B-C) Representative confocal immunofluorescent images showing SOX17-positive endodermal structures in ExEnd-aggregoids made of Flk1-GFP(B) or MP1 (*Bra*-GFP: *Sox17*-RFP) (C) cell lines. D) Circle plots demonstrating the ratio of ExEnd-aggregoids having SOX17-RFP-labeled ExEnd-like cells incorporated into the gut tube (“inside”). (E) Confocal immunofluorescent image showing total endoderm (immunostained SOX17) and the fluorescent signal from the Rosa26-BFP cell line used for obtaining RACL cells. (F) Representative phase contrast images with overlayed green (*Flk1*-GFP, from 2i-derived cells) and red (*Sox1*7-RFP, from ExEnd-derived cells) fluorescent signals of 120h NACLB- and XAL-aggregoids. G) Representative confocal immunofluorescent images of 120h NACLB- and XAL-aggregoids showing SOX17-RFP signal (from ExEnd-derived cells), FLK1-GFP signal (from 2i-derived cells), and immunofluorescence staining for SOX17. (H) UMAP projection of Sox17 expression across all conditions (control, RACL-, NACLB-, XAL-aggregoids) and time points. Sox17-positive clusters annotated as Endo-1 and Endo-2 are highlighted. (I) UMAP projection of *Ttr* (marker of ExEnd) and *Trh* (marker of DE) on Endo1 (DE) and Endo2 (ExEnd) clusters. (J) UMAP visualization of definitive and extraembryonic endoderm populations, illustrating cell type diversity across control, RACL-, NACLB-, and XAL-aggregoids (all time points). (K) Integrated UMAP of RACL-aggregoids (48–120 h) with projected pseudotime trajectories (Monocle3). The definitive endoderm cluster is highlighted. Extraembryonic endoderm populations were excluded from this analysis. The magnified fragment of the UMAP shows a region with annotated clusters (legend shared with panel J). (L) UMAP feature plots showing expression of selected genes in the highlighted region. (M) Dot plot showing expression of definitive and extraembryonic endoderm marker genes across annotated endodermal cell types in RACL-aggregoids. (N-P) Representative confocal immunofluorescent images showing the differences in endoderm organization between control gastruloids and RACL-aggregoid (O-P) at 96h (N-O) and 120h (P). SOX17 and BRA from reporter experssion, CDH1 from immunostaining. Q) Representative confocal immunofluorescent images of CDH1 and DAB2 expression in 120 h NACLB-aggregoids. (R-T) Chord plots based on CellChat showing the signaling relations between ExEnd and DE cell types.

Co-aggregation with ExEnd-like cells rescued this phenotype, resulting in the appearance of SOX17-positive cells, the formation of a tube-like structure (Figure 6B-C) and the emergence of two endodermal populations corresponding to ExEnd and DE (Figure 6H). Lineage tracing using ExEnd-like cells with *Sox17*-RFP or *Rosa26*-BFP reporters revealed distinct contributions of ExEnd cells. In RACL-aggregoids, ExEnd cells remained predominantly surface-localized, while the gut tube was largely derived from gastruloid-originating DE cells (Figure 6D, E). In contrast, NACLB- and XAL-aggregoids showed substantial incorporation of ExEnd cells into the gut epithelium (Figure 6D, F, G).

While both ExEnd and DE populations shared pan-endodermal markers such as *Sox17*, *Foxa2, Cdh1*, and *Gata6* (Figure 6M), they could be distinguished by the lineage-specific markers *Ttr*, *Amn, Apoe, Gata4, Pdgfra* and *Dab2* for ExEnd and *Foxa1*, *Trh, Shh, Car4* and *Kitl* for DE (Figure 6I, M). Clusters corresponding to ExEnd contained cells annotated as ParE and VE (Figure 6J). DE populations in RACL- and NACLB-aggregoids comprised multiple gut-associated lineages, including foregut, midgut, hindgut, and gut tube identities, whereas XAL-aggregoids were largely restricted to foregut-like states (Figure 6J). A subset of cells annotated as midgut was also detected within the ExEnd cluster across conditions, - and consistent with previous reports suggesting contributions of ExEnd to gut development(Kwon et al., 2008) (Figure 6J). Trajectory inference of RACL-aggregoids revealed a developmental progression from the PS (*T/Bra, Eomes)* branching to NM (*Mesp1, Cdh2*) and APS (*Foxa2, Gsc, Otx2*), followed by differentiation towards gut lineages (Figure 6K-M, Figure 5J).

A distinctive feature of ExEnd-aggregoids was the spatial organization of the emerging endoderm. In the absence of an extraembryonic scaffold, SOX17-expressing cells were located adjacent to the posterior BRA domain and formed dispersed clusters that could further merge(Hashmi et al., 2022; Vianello and Lutolf, 2021) (Figure 2I, 6N). In contrast, RACL- and NACLB-aggregoids displayed organized SOX17-positive domains flanking the midline and anterior BRA-positive regions (or only midline BRA domain in XAL-aggregoids) and forming a CDH1-positive ventral structure reminiscent of a notochordal plate(Balmer et al., 2016) (Figure 6O-P, Figure 5D, I). Notably, RACL- and NACLB-aggregoids showed a close spatial association between ExEnd and DE populations (Movie 03, Figure 6Q). CellChat analysis predicted multiple ligand–receptor interactions between ExEnd and DE populations, including LAMININ, BMP, non-canonical WNT, EPHA/B, and HH signaling (predominantly from ExEnd), as well as FN1 and VISFATIN (from DE) pathways, suggesting coordinated cross-regulation between these compartments (Figure 6R-T).

Together, these findings suggest that in aggregoids, ExEnd-like cells promote gut formation and stratification both by directly contributing to endodermal tissues and by enhancing endogenous DE specification and organization.

### ExEnd-aggregoids demonstrate enhanced mesoderm diversity

ExEnd-aggregoids exhibited increased abundance and diversity of mesodermal cell types compared to control gastruloids (Figure 7A). In controls, sparce mesoderm was largely restricted to paraxial derivatives, with somites primarily represented by dermomyotome-like populations (*Pax3, Dmrt2, Meox1, Tcf15*). In contrast, ExEnd-aggregoids contained additional ventral somitic derivatives, including sclerotome (*Pax1*) and endotome (*Kdr, Meox1*) (Figure 7A, B). Cell types predicted as IM and kidney primordium displayed limited expression of canonical IM markers such as *Osr1, Pax8* and *Pax2,* suggesting partial or immature identity (Figure 7B). Additionally, ExEnd-aggregoids contained cell types underrepresented in control gastruloids including axial mesoderm (*Noto, Shh, Foxa2, Chrd, Nog*), cardiopharyngeal mesoderm (*Tbx1, Isl1, Gata4, Tbx18, Tbx5, Hcn4, Prrx1* and *Prrx2*), and two endothelial clusters (*Nr2f2, Flt4, Kdr, Cdh5, Pecam1*) (Figure 7B, C).

**Figure 7.**
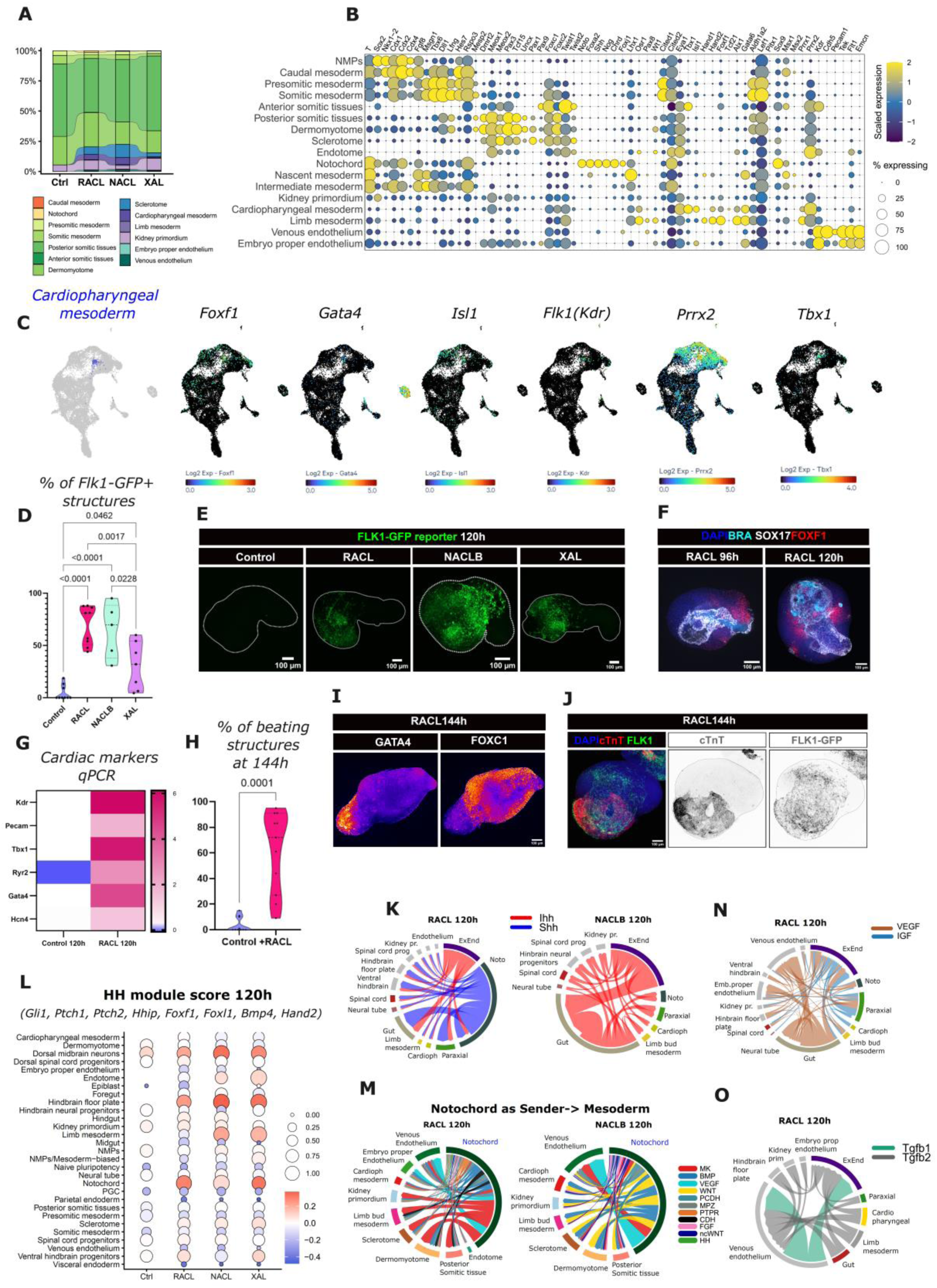
ExEnd-aggregoids demonstrate enhanced mesoderm diversity. (A) Alluvial plot showing proportions of mesodermal cell types within the total mesoderm population across conditions. (B) Dot plot showing expression of marker genes across mesodermal cell types. (C) UMAP projection of the cardiopharyngeal mesoderm cluster and its marker gene expression. (D) Violin plot showing percentage of structures with detectable FLK1-GFP signal at 120h per plate across conditions. Kruskal-Wallis test (n(control)=9, n(RACL)=9, n(NACLB)=5, n(XAL)=7). (E) Representative confocal fluorescent images showing *Flk1-*GFP reporter signal at 120h across conditions. (F) Representative confocal immunofluorescent images showing BRA, SOX17 and FOXF1 at 96-120h (G) Heatmap of qPCR analysis from pooled control gastruloids and RACL-aggregoids showing relative expression of cardiovascular marker genes at 120h (n=3, N=3). (H) Violin plot showing the proportion of structures exhibiting spontaneous beating activity at 144h in control gastruloids (n=10 biological replications) and RACL-aggregoids (n=11 biological replications). Statistical significance was assessed using non-parametric t test with Welch’s correction. (I-J) Representative confocal immunofluorescent images of 144h RACL-aggregoids. (K, N, M, O) Chord plots based on CellChat analysis showing the communication pairs with focus on HH signaling (K), notochord-derived signals (M) and crosstalk between gut and mesoderm (N, O).

The expansion of lateral plate mesoderm (LPM) and cardiovascular-associated lineages was particularly evident in RACL- and NACLB-aggregoids. *Flk1*(*Kdr*)-GFP signal, marking LPM and its derivatives, was detected in only 4,6% of control gastruloids at 120h, but increased to 69,65% of RACL-, 64,54% of NACLB-, and 30,73% of XAL-aggregoids (Figure 7D, E). Another LPM marker FOXF1 was also widely present in ExEnd-aggregoids but not in controls (Figure 7F). Cardiac and endothelial markers were detected by immunostaining (cTnT, GATA4) and qPCR (*Kdr, Pecam, Tbx1, Ryr2, Hcn4, Gata4*) (Figure7G, I, J). Spontaneous contractile activity emerged in 61,45% of RACL-aggregoids by 144 h compared to 3,7% in controls (Figure 7H; Movie 04). These data indicate that ExEnd co-aggregation was sufficient to enchance LPM formation and the initiation of downstream cardiovascular differentiation in the absence of exogenous cues(Rossi et al., 2021).

Mesoderm specification and patterning are known to depend on Hedgehog (HH) signaling(Borycki et al., 1998; Krauss et al., 1993; Newton et al., 2022; Prummel et al., 2020; Tsiairis and McMahon, 2009; Williams et al., 2010). CellChat(Jin et al., 2025) analysis revealed that at 120h the major source of HH ligands were ExEnd, notochord and gut endoderm - cell types which were represented in the ExEnd-aggregoids (especially in RACL- and NACLB-) but not in controls. In RACL-aggregoids, notochord-like cells acted as a source of SHH, with predicted interactions targeting paraxial mesoderm, neural tissues, and gut populations (Figure 7K), consistent with its patterning role in the embryo(Krauss et al., 1993; Stemple, 2005), HH module scores in the predicted clusters (Figure 7L) and the appearance of ventral-specific populations such as sclerotome and hindbrain floor plate. Importantly, HH signaling was not restricted to axial sources: RACL- and NACLB-aggregoids also expressed IHH, and in NACLB-conditions IHH signaling from both ExEnd and DE targeted IM, LPM, and somitic derivatives (Figure 7K). Besides HH, notochord-like cells acted as a major signaling source, exhibiting strong and diverse outgoing activity across multiple pathways, including MDK, BMP, VEGFA, and ECM-associated signaling with multiple mesodermal (endothelial, IM, cardioharyngeal cells, somitic tissues) targets (Figure 7M). In addition, we identified TGFb and VEGFa signaling interactions with gut populations as signaling sources and endothelial cells as receivers (Figure7N-O) consistent with the spatial proximity of *Flk1*-GFP positive endothelium to the gut in RACL- and NACLB-aggregoids (Figure 6B) as well as the association between FOXF1-positive cells and endoderm at 120h (Figure 7F).

Together, these results indicate that ExEnd co-aggregation establishes a coordinated signaling environment in which endodermal and axial compartments act as interacting signaling centers. Through combined HH and VEGF inputs, this niche promotes ventralization of mesoderm, supports LPM specification, and enables the emergence of cardiovascular lineages.

## DISCUSSION

Gastruloids represent SCBEMs generated from a single cell type (ESCs), requiring only minimal triggers, such as a transient CHIR pulse, to undergo a cascade of morphogenetic events recapitulating key aspects of mammalian gastrulation(Beccari et al., 2018; Brink and van Oudenaarden, 2021; Turner et al., 2017; Turner and Martinez Arias, 2024; van den Brink et al., 2014; Veenvliet et al., 2020; Veenvliet et al., 2021). Unlike blastoids(Jorgensen et al., 2026; Li et al., 2019; Rivron et al., 2018) or ETX embryos(Amadei et al., 2021; Lau et al., 2022; Sozen et al., 2018; Tarazi et al., 2022; Yilmaz et al., 2025), canonical gastruloids lack extraembryonic tissues(Arias et al., 2022; Liu and Warmflash, 2021) and hence develop in the absence of a spatially organized signaling architecture that those tissues provide during embryogenesis. However, the reductionist nature of gastruloids makes them suitable as a platform for modular engineering of developmental systems. For example, and as shown in our work, the controlled addition of specific embryonic and extraembryonic components can provide a framework for both dissecting developmental mechanisms that are not present in canonical gastruloids and for generating defined advanced morphotypes.

Here, we established a modular system to model the interactions between ExEnd and epiblast in mouse SCBEMs. Unlike XEGs, SCBEMs generated by co-aggregation with embryo-derived XEN cells(Berenger-Currias et al., 2022), ExEnd-aggregoids did not exhibit extensive neuralization. The obtained co-aggregated SCBEMs showed the emergence of DE, anterior and axial mesoderm, as well as a transcriptional signature enriched for NODAL targets, that are broadly consistent with previous reports demonstrating that Activin A treatment promotes a NODAL-driven developmental program in gastruloids(Arias et al., 2025; Dias et al., 2025; Wehmeyer et al., 2025). Recent studies have proposed that mammalian PS formation is orchestrated by two regulatory modules: an early/anterior module associated with EOMES and NODAL activity, and a late/posterior module that is dependent on WNT3A and T/BRA signaling(Arias et al., 2025; Dias et al., 2025; Schule et al., 2023; Wehmeyer et al., 2025). In contrast to CHIR-treated (canonical) and Activin A-treated gastruloids, both early/anterior and late late/posterior) PS programs were active in ExEnd-aggregoids resulting in a robust elongation and lineage diversity.

We did not find evidence that ExEnd-like cells act as TGFβ ligand senders in our model, nor did we detect substantial differences in the expression of NODAL-associated genes between control gastruloids and ExEnd-aggregoids prior to CHIR treatment. However, the significant upregulation of NODAL module following CHIR exposure in ExEnd-aggregoids but not in standard gastruloids might suggest the involvement of indirect mechanism based on response to CHIR. That is consistent with the effects of low dose of CHIR resulting in transcription profile similar to Activin A-treated gastruloids at 72h(Dias et al., 2025).

We propose that the partial capping of the epiblast by DKK1-expressing ExEnd-like cells lead to a gradient of differential response to the CHIR treatment, and a spatially compartmentalization of signal responsiveness within the aggregates. Notably, the addition of XAL cells resulted in fewer anterior mesoderm- and DE-enriched phenotypes compared with RACL- and NACLB-aggregoids, a difference that may be related to the comparatively low expression of *Dkk1* in XAL cells among the three ExEnd-like cell types tested. The proposed spatial compartmentalization was reflected by an initially asymmetric T/BRA expression at 72 h, emerging at the future posterior pole of the aggregate and a subsequent expansion towards the anterior pole, that broadly resembled the spatial dynamics of T/BRA expression associated with PS formation and extension in the mouse embryo(Rivera-Pérez and Magnuson, 2005).

The previously published mSCBEM protocol based on merging BMP4-treated and naïve aggregates, demonstrated similar dynamics of T/BRA distribution and the appearance of structures corresponding to node, notochord, blood vessels and cardiac mesoderm(Xu et al., 2021). Hence, despite relying on different experimental strategies, both the system of Xu et al. 2021(Xu et al., 2021) and our ExEnd co-aggregoids generated remarkably similar morphotypes. In Xu et al., 2021 patterning was driven by a localized signaling center, whereas in our aggregoids regionalization was likely achieved through regional modulation of morphogen activity by ExEnd-like cells. These findings suggest that early spatial compartmentalization of signaling cues may be sufficient to promote increased developmental complexity.

The presented modular approach, in our case a set of extraembryonic cell types with varying properties in combination with a canonical gastruloid, allows in principle to create a plethora of specific morphotypes, providing a strong argument for using co-aggregation as a versatile strategy for engineering SCBEMs.

## LIMITATIONS OF STUDY

Although gene expression analysis ExEnd-like populations revealed *Cer1* and *Lefty* expression in XAL cells and their resemblance to AVE, those markers were almost lost after co-aggregation and incubation in N2B27. This might be consistent with the transient expression of *Cer1* and *Lefty in vivo* and in embryo models(Nowotschin et al., 2019; Schumacher et al., 2024). As result, we did not observe the reproducible formation of rostral neuroectoderm upon co-aggregation with XAL cells, an outcome that would be expected from co-aggregation with cells possessing AVE-like identity(Andoniadou and Martinez-Barbera, 2013; Levine and Brivanlou, 2007). This problem might be approached by changing co-aggregation strategy: for example, adding XAL cells at the time point closer to CHIR pulse.

## ACKNOWLEDGMENTS

The project has received funding from the University of Oslo (UiO), the University Hospital of Oslo, the Research Council of Norway through its Centers of Excellence scheme grant Grant ID 262613, EU’s H2020 Marie Skłodowska-Curie Actions / COFUND Grant ID 801133, the European Innovation Council (EIC) (Horizon Europe) Pathfinder Challenge “Supervised Morphogenesis in Gastruloids” Grant ID 101071203 and the UIO:Life Science Convergence environments project ITOM. We thank Anna Bigas and Susanne van den Brink (Hospital del Mar Medical Research Institute, Barcelona, Spain) for providing *Rosa26-*BFP:*Flk1*-GFP mESC, Alexander Medvinsky (The University of Edinbugh, Edinburgh, UK) for providing *Flk1*-GFP mESC and Iftach Nachman (Tel-Aviv University, Israel) for providing *Bra*-GFP: *Sox17*-RFP mESC. We would like to thank Xian Hu (Edna) from the NorMIC Imaging Platform at the Department of Biosciences, University of Oslo, for providing assistances and access to the spinning disk microscope for imaging of gastruloids. Figures 1A and 2A were created with BioRender.com.

## AUTHOR CONTRIBUTIONS

N.P.S. and S.V.P. contributed equally. N.P.S., S.V.P. and S.K.D. conceived and designed the experiments; N.P.S., S.V.P. and B.K.C. carried out the experiments, N.P.S., S.V.P., J.V.V., T.M.L.A., M.L. and T.C. analyzed and elaborated data, T.M.L.A. and J.Ø. executed the scRNA-seq analysis; J.V.V. and E.M. provided critical comments to the manuscript; N.P.S., S.V.P. and S.K.D. wrote the manuscript; J.V.V. and S.K.D. provided funds.

## MATERIALS AND METHODS

### Mouse Pluripotent Stem Cell culture and naive conversion

The following mouse embryonic stem cell (mESC) lines were used in this study: R1 (SCRC-1011, ATCC), *Flk1-eGFP(Jakobsson et al., 2010)*, provided by Alexander Medvinsky’ laboratory *Rosa26-BFP::Flk1-GFP(Ragusa et al., 2025)*, provided by Anna Bigas’ laboratory), *and Bra-GFP/Sox17-RFP(Pour et al., 2022)*, provided by Iftach Nachman’s laboratory). mESCs were cultured on 6-well culture plates (140675, ThermoFisher) covered with 0,1% Gelatine solution (ES-006-B, Merck) in serum + LIF medium consisting of KnockOut DMEM (10829018, Gibco) supplemented with 15% fetal bovine serum (FBS; 10309433, HyClone), non-essential amino acids (NEAA; 11140050, Gibco), GlutaMAX (35050061, Gibco), 0.1 mM β-mercaptoethanol (M3148, Merck), and mouse leukemia inhibitory factor (LIF; ESG1107, Merck).

For naïve state conversion, mESCs were plated on wells coated with 0.01% ornithine (P3655, Merck) and cultured in 2i/N2B27+LIF medium. 2i/N2B27+LIF composition was: 1:1 mixture of DMEM/F12 (11330032 or 11039021, Gibco) and Neurobasal medium (21103049 or 12348017, Gibco), supplemented with N2 (17502048, Gibco), B27 (17504044, Gibco), GlutaMAX (35050061, Gibco), NEAA (11140050, Gibco), sodium pyruvate (11360039, Gibco), β-mercaptoethanol (M3148, Merck), bovine serum albumin (BSA; A8412, Merck), LIF (ESG1107, Merck), CHIR99021 (3 µM; 4423, Tocris), and PD0325901 (1 µM; 4192, Tocris). Cells were maintained in naïve conditions for 5–7 days prior to downstream applications.

All cell cultures were incubated at 37 °C in 100% humidity with 5% CO₂. Both standard and 2i/N2B27+LIF media were refreshed daily. Cells were passaged every other day using Accutase (A6964, Merck). All cell cultures were routinely tested for Mycoplasma using Mycoalert Mycoplasma Detection Kit (LT07-318, Lonza).

### Differentiation of ExEnd-like cell types

mESCs cultured in 2i/N2B27+LIF medium for 5–7 days were harvested using Accutase (A6964, Merck) or TrypLE Express (12604013, Gibco) and replated at a density of 1×10⁵ cells/cm² on 0.1% gelatin-coated 6-well plates. Cells were seeded in RPMI 1640 medium (11875093, Gibco) supplemented with 1×B27 Supplement Minus Insulin (A1895601, Gibco). After first 24 hours, the medium was replaced with RACL medium, consisting of RPMI 1640 (11875093, Gibco), 1×B27 Supplement Minus Insulin (A1895601, Gibco), 100 ng/mL ActivinA (Peprotech, 120-14P), 3 μM CHIR99021 (4423, Tocris Bioscience), and mouse LIF (ESG1107, Merck). The cultures were passaged once between days 3 and 4 of the protocol. On days 6–8 of RACL treatment, cells were either used for co-aggregation experiments in RACL conditions or subjected to further differentiation protocols.

To initiate further differentiation, RACL medium was replaced with NACL medium for 24 hours. NACL medium consisted of N2B27 basal medium composed of 1:1 mix of DMEM/F12 (11320033, Gibco) and Neurobasal medium (21103049, Gibco) supplemented with N2 Supplement (17502048, Gibco), B-27 Supplement (17504044, Gibco), GlutaMAX (35050061, Gibco), non-essential amino acids (11140050, Gibco), sodium pyruvate (11360039, Gibco), β-mercaptoethanol (M3148, Merck), and BSA (A8412, Merck), supplemented with 100 ng/mL Activin A (Peprotech, 120-14P), 3 μM CHIR99021 (4423, Tocris Bioscience), and mouse LIF (ESG1107, Merck).

For NACLB differentiation, NACL medium was further supplemented with 50 ng/mL recombinant murine BMP4 (Peprotech, 315-27). To generate anterior visceral endoderm-like (AVE-like) cells, cells previously maintained in NACL were treated with XAL medium, consisting of N2B27 supplemented with 10 μM XAV939 (S1180, Selleck Chemicals), 100 ng/mL Activin A (120-14P, Peprotech), and mouse LIF (ESG1107, Merck). Both NACL+B and XAL treatments were applied for three days.

### Generation of gastruloids and aggregoids

Gastruloids were generated following previously established protocols(Baillie-Johnson et al., 2015; Beccari et al., 2018). Briefly, 300 mouse embryonic stem cells (mESCs) were plated in 40 µL of NDiff 227 medium (Y40002, Takara) or home-made N2B27 medium (composition described above) into each well of a 96-well U-bottom cell-repellent microplate (650970, Greiner Bio-One). Naïve mESCs grown for 5-7 days in 2i/N2B27+LIF conditions were used as starting material. At 48 hours post-aggregation, 150 µL of NDiff or N2B27 medium supplemented with 3 µg/mL of CHIR99021 was added to each well. Media change was performed every 24 hours by removing 150 µL and replacing it with fresh NDiff or N2B27 medium. Gastruloids were harvested for analysis at 48, 72, 96, 120, 144, and 168 hours.

To generate aggregoids, 300 naïve mESCs were mixed with 100 additional cells (RACL, NACLB, or XAL) at the aggregation step. The cell mixture was plated under the same conditions as described above for gastruloid formation. All subsequent steps, including CHIR99021 treatment and daily media changes, were performed as described for the standard gastruloid protocol.

### Microscopy and live imaging

Imaging of living and fixed cell cultures was performed EVOS M7000 (Thermo Fisher Scientific) microscope. Life imaging and time lapse imaging of gastruloids and ExEnd-aggregoids with Phase Contrast, Brightfield, GFP, and RFP channels were made using IncuCyte S3 (Sartorius).

Confocal time lapse images of RACL-aggregoids with GFP, RFP, and BFP channels were acquired using Andor Dragonfly Spinning Disk confocal microscope (Dragonfly) with Zyla sCMOS camera, 25-micron pinhole disk and Nikon 20X/0.70 or 10X/0.40 air objectives. Images were taken every 15 minutes for a total duration of 24 hours. The microscope was equiped with a weather chamber from Okolab, which maintained 5% CO₂ and appropriate humidity levels throughout the imaging period. The Perfect Focus System (PFS) was utilized to ensure the focus of the sample was maintained during the entire imaging process.

Confocal imaging of fixed and immunostained gastruloids and ExEnd-aggregoids was done using Nikon Crest Spinning disk (X-light V3 CREST).

### Immunofluorescent staining and microscopy of fixed samples

Gastruloids and cell culture were fixed in 4% PFA for 1h at room temperature on a shaking platform then washed three times with PBS and stored at +4°C in PBS.

For immunostaining the specimens were first incubated in blotting solution: PBS (14190, Gibco), 10% BSA (422361V, VWR Life Science), 0,5% Triton x-100 (M143, VWR Life Science), 0,02 % SDS (L3771, Sigma) for 1h at the room temperature, then incubated with first antibodies diluted in the blotting solution for overnight at +4°C. After the incubation with primary antibodies the gastruloids were washed with blotting solution 3 times for 30 min at the room temperature and incubated with secondary antibodies diluted in blotting solution for 3h at the room temperature. After the incubation with secondary antibodies the gastruloids were washed with blotting solution 3 times for 30 min.

The following primary antibodies were used: goat anti-Brachyury (1:200, R&D Systems, AF2085), goat anti-Cdh1 (E-cadherin) (1:200, R&D Systems, AF748), rabbit anti-Cdx2 (1:200, ThermoFisher, MA5-14494), mouse anti-Cardiac Troponin T (1:200, ThermoFisher,MA5-12960), mouse anti-Dab2 (1:200, Santa Cruz, sc-136964), mouse anti-Dkk1 (1:100, Santa Cruz, sc-374574), rat anti-Eomesodermin (1:100, ThermoFisher, 14-4875-82), rabbit anti-FoxA2 (1:200, Merck-Millipore, 07-633), rabbit anti-FoxC1 (1:200, Abcam, ab223850), rabbit anti-FoxF1 (1:200, Abcam, ab308633), rabbit anti-FoxJ1 (1:200, Abcam, ab235445), mouse anti-Gata4 (1:200, Santa Cruz, sc-25310), rabbit anti-Lefty (1:100, ThermoFisher, BS-11236R), rabbit anti-Otx2 (1:200, ThermoFisher, 13497-1-AP), rat anti-PDGFRα (1:200, ThermoFisher, 14-1401-82), mouse anti-Sox2 (1:200, R&D Systems, MAB2018), rabbit anti-Sox17 (1:200, Abcam, ab224637).

The following secondary antibodies were used: donkey anti-rabbit Cy3 (1:500, Jackson ImmunoResearch, 711-165-152), donkey anti-rabbit AlexaFluor 647 (1:500, Jackson ImmunoResearch, 711-605-152), donkey anti-mouse Cy3 (1:500, Jackson ImmunoResearch, 715-165-150), donkey anti-mouse AlexaFluor 488 (1:500, Jackson ImmunoResearch, 715-545-150), donkey anti-mouse AlexaFluor 647 (1:500, Jackson ImmunoResearch, 715-605-150), donkey anti-goat Cy3 (1:500, Jackson ImmunoResearch, 705-165-003), donkey anti-goat AlexaFluor 488 (1:500, Jackson ImmunoResearch, 705-545-147), donkey anti-goat AlexaFluor 647 (1:500, ThermoFisher, A32849), donkey anti-rat Cy3 (1:500, Jackson ImmunoResearch, 712-165-153), donkey anti-rat AlexaFluor 488 (1:500, ThermoFisher, A48269).

All the incubations and washing steps were performed at the shaking conditions. After the staining the gastruloids were mounted in a RapiClear solution (152001, SunJin Lab) between slide and cover slip. The samples were visualized with X Light V3 CREST confocal spinning disc microscope (Nikon).

### Image analysis

Images were processed using Fiji (ImageJ), an open-source image analysis software(Schindelin et al., 2012) Dotted line pluggin was used for drawing image outline (https://imagej.net/ij/plugins/dotted-line.html).

Midline lengths, area, and GFP and RFP intensity measurements of the gastruloids were done using an in-house developed script coded with Python3. Brightfield images were segmented using a two-steps algorithm. First, the gastruloid was roughly segmented using a gradient-boosting classifier (CatBoost)(Prokhorenkova et al., 2018), using edges and textures averaged at different scales (using skimage’s ‘multiscale_basic_features’(van der Walt et al., 2014)) as features and trained on a few hand-segmented images with the threshold chosen to maximise the F1-score. Secondly, the center of gravity of the biggest object segmented at this point was fed as an input point for the Segment Anything Model (SAM)(Kirillov et al., 2023), resulting in an high-quality mask of the gastruloid. If the mask was not touching any edge of the image, otherwise the image would not be considered for analysis, the midline of the gastruloid was computed using an algorithm adapted from MOrgAna(Gritti et al., 2021), and the mask was divided into 10 sections spanning an equal amount of the midline. For each of these sections, the size and the fluorescence intensities, extracted from the corresponding fluorescence images, were computed and saved as a text file for final analysis. Elongation index was calculated as the ratio of gastruloid midline length to the square root of projected area:

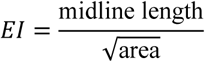

### RNA extraction and RT-qPCR

Total RNA was extracted from either cell cultures or pooled gastruloids using the RNeasy Micro Kit (Qiagen, 74004), following the manufacturer’s instructions. For each extraction, 48–96 gastruloids or 300,000–1,000,000 cells were collected and stored at –80 °C until processing. RNA concentration was quantified using a Nanodrop spectrophotometer (ThermoFisher Scientific).

Complementary DNA (cDNA) was synthesized using the High-Capacity cDNA Reverse Transcription Kit (ThermoFisher Scientific, 4368814). Quantitative real-time PCR (RT-qPCR) was performed using a ViiA 7 thermocycler (ThermoFisher Scientific) with TaqMan Gene Expression Master Mix and gene-specific TaqMan probes (ThermoFisher Scientific).

Each qPCR reaction contained 1 μL of cDNA (diluted to 20 ng/μL) in a total volume of 10 μL, conducted in 386-well plates (4309849, Applied Biosystems). Gene expression levels were normalized to the housekeeping gene *Gapdh*, and relative expression was calculated using the 2^–ΔΔCT method. The number of technical replications was 3, the number of biological replications was a minimum 3. 2i/N2B27 cells were used as a reference sample for normalization.

Primer details are listed below:

|  |  |  |
| --- | --- | --- |
| TaqMan gene expression assay Mm00515474_m1 (Cer1) | ThermoFisher | Cat# 4331182 |
| TaqMan gene expression assay Mm00438422_m1 (Dkk1) | ThermoFisher | Cat# 4331182 |
| TaqMan gene expression assay Mm00484689_m1 (Gata4) | ThermoFisher | Cat# 4453320 |
| TaqMan gene expression assay Mm00802636_m1 (Gata6) | ThermoFisher | Cat# 4331182 |
| TaqMan gene expression assay Mm01176086_m1 (Hcn4) | ThermoFisher | Cat# 4331182 |
| TaqMan gene expression assay Mm00433954_m1 (Hhex) | ThermoFisher | Cat# 4331182 |
| TaqMan gene expression assay Mm01247712_m1 (Hnf4a) | ThermoFisher | Cat# 4453320 |
| TaqMan gene expression assay Mm00439613_m1 (Ihh) | ThermoFisher | Cat# 4331182 |
| TaqMan gene expression assay Mm01222421_m1 (Kdr1) | ThermoFisher | Cat# 4331182 |
| TaqMan gene expression assay Mm01226102_m1 (Lama1) | ThermoFisher | Cat# 4448892 |
| TaqMan gene expression assay Mm03053915_s1 (Lefty1) | ThermoFisher | Cat# 4331182 |
| TaqMan gene expression assay Mm00446859_m1 (Otx2) | ThermoFisher | Cat# 4331182 |
| TaqMan gene expression assay Mm00440701_m1 (Pdgfra) | ThermoFisher | Cat# 4448892 |
| TaqMan gene expression assay Mm01242576_m1 (Pecam1) | ThermoFisher | Cat# 4448892 |
| TaqMan gene expression assay Mm00465877_m1 (Ryr2) | ThermoFisher | Cat# 4331182 |
| TaqMan gene expression assay Mm00488363_m1 (Sox17) | ThermoFisher | Cat# 4331183 |
| TaqMan gene expression assay Mm00448949_m1 (Tbx1) | ThermoFisher | Cat# 4331182 |
| TaqMan gene expression assay Mm01180221_g1 (Gapdh) | ThermoFisher | Cat# 4331182 |

### Bulk RNA-sequencing

Total RNA was extracted from totally 12 mouse cell samples: naive mESCs in 2i/N2B27 conditions and differentiated extraembryonic endoderm-like cell types: RACL, NACLB, and XAL. 2 µg of total RNA in 100ul of RNAse-free water was used for library preparation. RNA quality was assessed using an Agilent Bioanalyzer. RNA-seq libraries were prepared using a strand-specific, poly(A)-selected protocol, TruSeq Stranded mRNA Library Prep (20020595, Illumina), and adapters TruSeq RNA UD Indexes (2002237, Illumina). Sequencing was performed on an Illumina NovaSeq X Plus system using 1.5B flow cells, generating paired-end 150 bp reads. Libraries were multiplexed and sequenced across two lanes, and FASTQ files from the two lanes were merged prior to downstream analysis. Sequencing instrumentation and software: NovaSeqXPlus System Suite v1.2.2.48004, BCL Convert v4.1.23, NSC scripts v1.0. Adapter and quality trimming was performed using BBDuk (BBMap toolkit v34.56), which was also used to remove PhiX spike-in sequences. BBDuk was run with adapter trimming from the 3′ end using k-mer matching (k=23, mink=11, hdist=1), quality trimming at a Phred threshold of 15 (qtrim=r, trimq=15), and reads were discarded if their average quality fell below 15 or their length dropped under 36 bp, with a hard right-clip applied at position 150.

The resulting high-quality reads were aligned to the mouse reference genome (Mus_musculus.GRCm39.dna.toplevel.fa) and transcriptome (Mus_musculus.GRCm39.113.gtf) using HISAT2 (v2.1.0)(Kim et al., 2019) with default parameters, setting the strandedness to RF. Gene-level read counts were subsequently quantified using featureCounts (v1.4.6-p1)(Liao et al., 2019) with paired-end and reverse-strand options enabled (-p -s 2).

### Differential expression analysis of bulk RNA sequencing data

The gene counts from the featureCounts output were loaded into R (v4.5.1) and combined across all samples. A preliminary quality check was conducted to identify any technical outliers or anomalies. Genes were filtered out with fewer than 50 total counts across the samples, ensuring they were expressed in at least two of the eight samples.

For exploratory analysis, VST-normalized counts were Z-score transformed and the top 500 most variable genes were selected based on variance across samples. Principal component analysis (PCA) was performed on these genes, and the genes with the highest loadings for PC1 and PC2 were identified (prcomp). Sample-level relationships were further assessed by hierarchical clustering using Euclidean distance with Ward’s D2 linkage (hclust), and by sample-to-sample correlation heatmaps generated using pheatmap (v1.0.13).

The quality-controlled count matrix was loaded into DESeq2 (v1.48.1)(Love et al., 2014) using condition as the design variable (2i, RACL, NACLB and XAL), with size factors estimated using the ratio method. Differentially expressed genes were identified using the Wald test across all pairwise condition comparisons, with genes meeting an adjusted p-value threshold of 0.05 and an absolute log2 fold change greater than 2 considered significant. MA and volcano plots were generated for each comparison. Gene over-representation and pathway analyses were performed on up-and down-regulated genes using clusterProfiler (v4.16.0)(Wu et al., 2021). GO biological process (BP) enrichment was performed using enrichGO with gene symbols mapped against org.Mm.eg.db (v3.12.0), and KEGG pathway enrichment was performed using enrichKEGG with gene symbols first converted to Entrez IDs via bitr function, using the mouse KEGG organism code (mmu). In both cases, Benjamini-Hochberg adjusted p-values were used for multiple testing correction, with a significance threshold of 0.05.

### Bulk RNA-seq marker-gene visualisation

Filtered gene-level bulk RNA-seq counts (three replicates per condition) were library-size normalised to counts per million and log2-transformed (log2[CPM + 1]) using edgeR. Expression of a curated panel of lineage marker genes was displayed as a heatmap (pheatmap), with each gene scaled to a z-score across samples to show relative expression. Genes were ordered by lineage and grouped into blocks comprised of representative markers for naive pluripotency/epiblast, endoderm, primitive endoderm, anterior visceral endoderm, parietal endoderm, and extraembryonic visceral endoderm.

### Integration of scRNA-seq datasets for pseudobulk comparison to bulk RNA data

Raw count data from five published mouse embryo scRNA-seq datasets(Argelaguet et al., 2019; Cheng et al., 2019; Liu et al., 2022; Mohammed et al., 2017; Thowfeequ et al., 2024) were integrated using Seurat v4(Hao et al., 2021). For datasets where genes were represented as Ensembl IDs(Argelaguet et al., 2019; Thowfeequ et al., 2024), gene identifiers were converted to HGNC symbols using biomaRt(Durinck et al., 2009). Low-quality cells were removed by retaining only those with over 5,000 detected features (genes). Each dataset was independently normalised using Seurat’s default log-normalisation and the top 2,000 highly variable genes identified using the variance-stabilising transformation (VST) method. Datasets were integrated using canonical correlation analysis (CCA)-based anchor finding (FindIntegrationAnchors / IntegrateData), and the integrated assay was used for downstream dimensionality reduction. Principal component analysis (PCA) was performed on the top 50 PCs, and cells were embedded in two dimensions using UMAP based on the first 30 PCs. Cell type identities were assigned to each cell by transferring lineage annotations from the original metadata of each dataset, which were then manually curated and refined using marker gene expression.

To enable comparison with bulk RNA-seq data from *in vitro*-differentiated endoderm cells, pseudobulk expression profiles were generated for each annotated cell type by summing raw counts across all cells of a given type (AggregateExpression). Pseudobulk and bulk RNA-seq count matrices were combined on their common gene set and library-size normalised to counts per million (log-CPM) using edgeR(Chen et al., 2025). To mitigate systematic differences between data modalities, batch correction was applied using ComBat (sva package)(Leek et al., 2012), with pseudobulk and bulk samples treated as separate batches. A correlation matrix showing the similarity between *in vitro* bulk RNA-seq samples and *in vivo* scRNA-seq pseudobulk profiles was assessed by Pearson correlation and visualised as heatmaps (pheatmap). For this analysis, the gene set was restricted to the top 100 highly variable genes across samples, with cell-cycle-associated and housekeeping genes excluded. Principal component analysis was assessed using the top 5000 highly variable genes across samples to visualise the global relationship between pseudobulk and bulk RNA-seq samples.

### Single cell suspension preparation and fixation

Gastruloids corresponding to four conditions: standard gastruloid, RACL-aggregoids, NACLB- aggregoids, XAL-aggregoids, - were collected at 48h, 72h, 96h, and 120h. For each condition: 384 gastruloids were pooled at 48h, 288 gastruloids at 72h, 192 gastruloids at 96h and 96 gastruloids at 120h. For preparation of single cell suspension, the gastruloids were dissociated according to published protocol(Bolondi et al., 2021) with our modifications. The objects were collected and transferred to a Petri dish with 200 µl of TrypLE Express (12604013, Gibco), incubated at 37°C with pipetting every 5 min until full dissociation to a single-cell suspension. Then TryLE was quenched with 800 µl of ice-cold PBS + 0,5% BSA (A8412, Merck). After filtering through a cell strainer (27215, Stemcell Technologies), the cell suspension was centrifuged at 4°C, 300 rcf for 5 min. After discarding the supernatant, the cell pellet was washed with 1 ml of PBS + 0,04% BSA and centrifuged at 4°C, 300 rcf for 5 min. Washing was repeated one more time and finally the cell pellet was dissociated in 1 ml of PBS + 0,04% BSA. Cell viability was assessed by Trypan Blue (15250-061, Gibco) staining. Number of dead cells was less than 5% for all stages and conditions. Then cell suspension was fixed using Chromium Next GEM Single Cell Fixed RNA Sample Preparation Kit, 16 rxns (1000414; 10X Genomics) and stored at −80C or +4C until library preparation. Samples with not less than 300 000 cells after fixation and washing were used for library construction.

### Library preparation and single cell RNA sequencing

Samples were multiplexed, and cDNA libraries construction was performed from 10 000 cells per sample with Chromium Fixed RNA Kit (1000496,10X Genomics) according to the user guide. Quality control and concentration measurements of cDNA libraries prior sequencing were made using Agilent BioAnalyzer high sensitivity DNA kit on Agilent 2100 Bioanalizer (Agilent Technologies). Single-cell RNA sequencing was done in Norwegian Sequencing Centre on NovaSeqXPlus (NovaSeqXPlus System Suite: 1.2.2.48004) platform 1 lane 25B 300 cycles.

### scRNA-seq processing and data analysis

The sequenced data was aligned against the murine mm10 reference genome using the 10x Genomic Cell Ranger (v8.0.1) pipeline, de-multiplexing the samples *in silico* accordingly. The cell barcode matrices generated by the CellRanger pipeline (v8.0.1) were loaded into R (v4.5.1) using the Seurat library (v5.3.0)(Hao et al., 2024). The first-pass quality control required cells to have at least 200 features and no more than 7500, and UMI counts to be between 400 and 40000. The mitochondrial gene content was set to below 25%. The mitochondrial gene content cut-off was determined by a data-driven method. Then, a standard Seurat pipeline was run to identify clusters, primarily based on the number of UMI and feature counts. Next, second-pass QC was performed by removing clusters mainly driven by low UMI/feature counts. We verified that removed clusters with low UMI/feature counts were solely due to low counts and not driven by biologically relevant signals, using Seurat’s FindMarker function.

After the initial quality control, cells were scored for doublets using scDblFinder (v1.22.0), and cells with a doublet score exceeding 0.7 were excluded from further analysis.

The quality-controlled cells were integrated based on condition (medium used) and time using Seurat’s RPCA-based integration method with 5000 integration features, after which the integrated object was scaled with cell cycle scores (S and G2M) regressed out, followed by PCA, UMAP, and clustering at resolution 0.5.

Further data analysis and visualizations were performed using R (v4.5.1 and 4.5.2) and Loupe browser.

### Integration

The quality-controlled cells were integrated based on conditions (medium used) and time using Seurat’s RPCA-based integration method. We used 5000 integration features for the process. Cells were annotated by transferring labels from the reference mouse embryo E6.5-E9.5 atlas(Pijuan-Sala et al., 2019). Some cell types were merged into a standard label to reflect the biological type.

### $Reference mapping

Cells were annotated by transferring labels from the Mouse gastrulation reference atlas (https://marionilab.github.io/ExtendedMouseAtlas/) using Seurat’s FindTransferAnchors and TransferData functions(Stuart et al., 2019) with PCA reduction and 30 dimensions, transferring the celltype_extended_atlas labels from the reference, and only predictions with a confidence score above 0.7 were retained.

The transferred labels were then manually curated per time point. Related cell types were merged into biologically meaningful groups: all cardiopharyngeal progenitor and cardiomyocyte subtypes (FHF and SHF variants) together with pharyngeal mesoderm were collapsed into “Cardiopharyngeal mesoderm”, and Foregut, Pharyngeal endoderm, and Thyroid primordium were merged into “Foregut”. Cells labeled as PGC were reassigned to “Naive pluripotency” if they co-expressed all four naive pluripotency markers (*Nanog, Zfp42, Klf2, Klf4*) and retained as PGC otherwise. Finally, cell types not expected in gastrulation were removed from the dataset before downstream analysis.

### Pseudotime analysis

Pseudotime trajectories were inferred per condition using Monocle3 (v1.4.26)(Trapnell et al., 2014). The manually curated Seurat objects were converted to CellDataSet objects, clustered, and fit with a principal graph using learn_graph (with use_partition = FALSE to learn a single graph across partitions). For non-control conditions, extraembryonic and visceral endoderm populations (Visceral endoderm, ExE endoderm, and Parietal endoderm; Parietal endoderm and Visceral endoderm for XAL) were removed prior to trajectory learning to restrict the analysis to embryonic lineages. Root cells were selected at the earliest time point (48h) as cells co-expressing the naive pluripotency markers *Nanog, Zfp42, Klf2*, and Klf4 while lacking expression of the differentiation markers *Dazl, Tfap2c, Fgf5*, and *Gata2*, and with a Marioni transfer confidence score above 0.7; for the control condition, a specific manually selected cell meeting these criteria was used directly (Ctrl72h_CGTTTGTCAGGATGCTACTTTAGG-1). Cells were then ordered along the principal graph with “order_cells” using the selected root, and the resulting pseudotime values were added to the cell metadata for downstream visualization and comparison across time points and cell types.

### Integration of gastruloids with mouse embryo reference

For integration(Stuart et al., 2019) of gastruloids with mouse embryo reference, the Extended Mouse Atlas(Imaz-Rosshandler et al., 2024; Pijuan-Sala et al., 2019) served as the in vivo reference. The published SingleCellExperiment 1.32.0 object(Amezquita et al., 2020) was imported into R (version 4.5.2) and converted to a Seurat (v5)(Hao et al., 2024) object retaining raw counts. Gene identifiers were converted from Ensembl IDs to MGI symbols using the atlas gene metadata, with duplicate symbols made unique. Per-cell quality metrics (genes and UMIs detected) were recomputed from the count matrix, and cells with fewer than 3,000 detected genes were removed. To prevent highly abundant cell types from dominating the integration, cells were randomly downsampled to a maximum of 2,000 per annotated cell type (celltype_extended_atlas) using a fixed random seed (42). Developmental stage (E6.5–E9.5) and cell-type annotations were retained from the original metadata.

Gastruloid libraries from four culture conditions (CTRL, RACL, NACLB, XAL), each sampled at [48, 72, 96 and 120 h], were imported as raw count matrices with associated metadata. Each condition was assembled into a Seurat object, labelled by condition of origin, and filtered to retain cells with at least 2,000 detected genes. The reference was held to a stricter gene-detection threshold than the query to ensure high-quality anchors for label transfer, while the more permissive cut-off applied to the gastruloid data preserved cell numbers in the query. Data integration. The downsampled reference and the four gastruloid condition datasets were assembled into a single object; only reference cells were down sampled (to a maximum of 2,000 per annotated cell type), while all gastruloid cells were retained. The combined object was split by sample of origin, and each dataset was independently log-normalized, with its 3,000 most variable features identified. A shared set of 3,000 integration features was selected across datasets, and each dataset was scaled and reduced by PCA on these features (40 components). Integration anchors were identified across all five batches (the reference atlas and the four gastruloid conditions) by reciprocal PCA (FindIntegrationAnchors, reduction = “rpca”, 40 dimensions) and used to compute a batch-corrected expression matrix (IntegrateData, 40 dimensions; Stuart et al., 2019). The integrated assay was scaled and reduced by PCA (50 components); the first 40 components were used to compute the UMAP embedding, a shared-nearest-neighbor graph. This integrated embedding was used for all presented analyses and visualization.

To characterize reference cell types within each developmental window, the atlas was subset by stage, and, within each stage, markers were identified for every cell type represented by at least 10 cells (FindAllMarkers on log-normalized RNA-assay values; Wilcoxon rank-sum test; positive markers only; minimum 25% detection, ≥10% detection difference, ≥0.25 log2 fold-change). Markers with a Benjamini–Hochberg adjusted p < 0.05 were retained. UMAP projections were generated in ggplot2 (v4.0.3) with point layers rasterized at 600 dpi (ggrastr 1.0.2); cell types were colored with a lineage-structured palette and labelled at cluster centroids.

### Cell–cell communication analysis

Cell–cell communication was inferred per condition using CellChat (v2.2.0)(Jin et al., 2025) with the mouse CellChatDB as the ligand–receptor reference. For each condition, the manually curated Seurat object was loaded, and cell type identities were set from the manual_marioni_label column. Cell types with fewer than 10 cells were excluded to stabilize the probability estimates. Log-normalized expression from the RNA assay and the corresponding cell type labels were used to construct the CellChat object. The standard CellChat pipeline was then run to identify ligand–receptor interactions per cell group, and communication probabilities were computed at both the interaction and pathway level using the triMean method. Finally, the networks were aggregated and centrality scored via netAnalysis_computeCentrality, and the resulting communication pairs and pathway-level networks were used for downstream comparison across conditions.

### Calculation of module scores

Pathway-associated transcriptional module scores were computed using Seurat AddModuleScore fuction. Gene sets for signaling pathways were following: *Lefty1, Lefty2, Smad7, Tgif1, Skil, Pitx2, Eomes, Foxa2, Gsc, Cer1, Lhx1, Mixl1, Otx*2 for Nodal module; *Axin2, Lef1, Sp5, Notum, Tcf7, Cdx1, Cdx2, Cdx4, Msgn1, Tbx6, Dkk1, Ccnd1* for Wnt module; *Id1, Id2, Id3, Smad6, Smad7, Msx1, Msx2, Bmp4, Bmp7, Hand1* for Bmp module; *Dusp6, Spry1, Spry2, Etv4, Etv5, Fgf8, Fgf4, Cdx4, Msgn1* for Fgf module; *Gli1, Ptch1, Ptch2, Hhip, Foxf1, Foxl1, Bmp4, Hand2* for Hh module; *Cyp26a1, Cyp26b1, Rarb, Rara, Crabp1, Crabp2, Hoxa1, Hoxb1* for RA module, *Hes1, Hey1, Lfng, Dll1, Dll3, Notch1, Notch2, Nrarp* for Notch module.

### Statistical analysis

Statistical analyses were performed using GraphPad Prism 10 (Version 10.2.0). Data are presented as mean ± SEM unless otherwise indicated. Individual data points represent biological replicates or individual structures. For comparisons involving multiple groups, Kruskal-Wallis tests with Dunn’s multiple comparisons tests were used. For comparisons involving two groups, two-tailed unpaired Welch’s t-tests were used. Asterisks indicate significance level: ns *p* ≥ 0.05, ∗ *p* ≤ 0.05, ∗∗ *p* ≤ 0.01, ∗∗∗ *p* ≤ 0.001, ∗∗∗∗ *p* ≤ 0.0001. Difference was referred to as significant when *p* ≤ 0.05.

---

**Supplementary figure 1.**
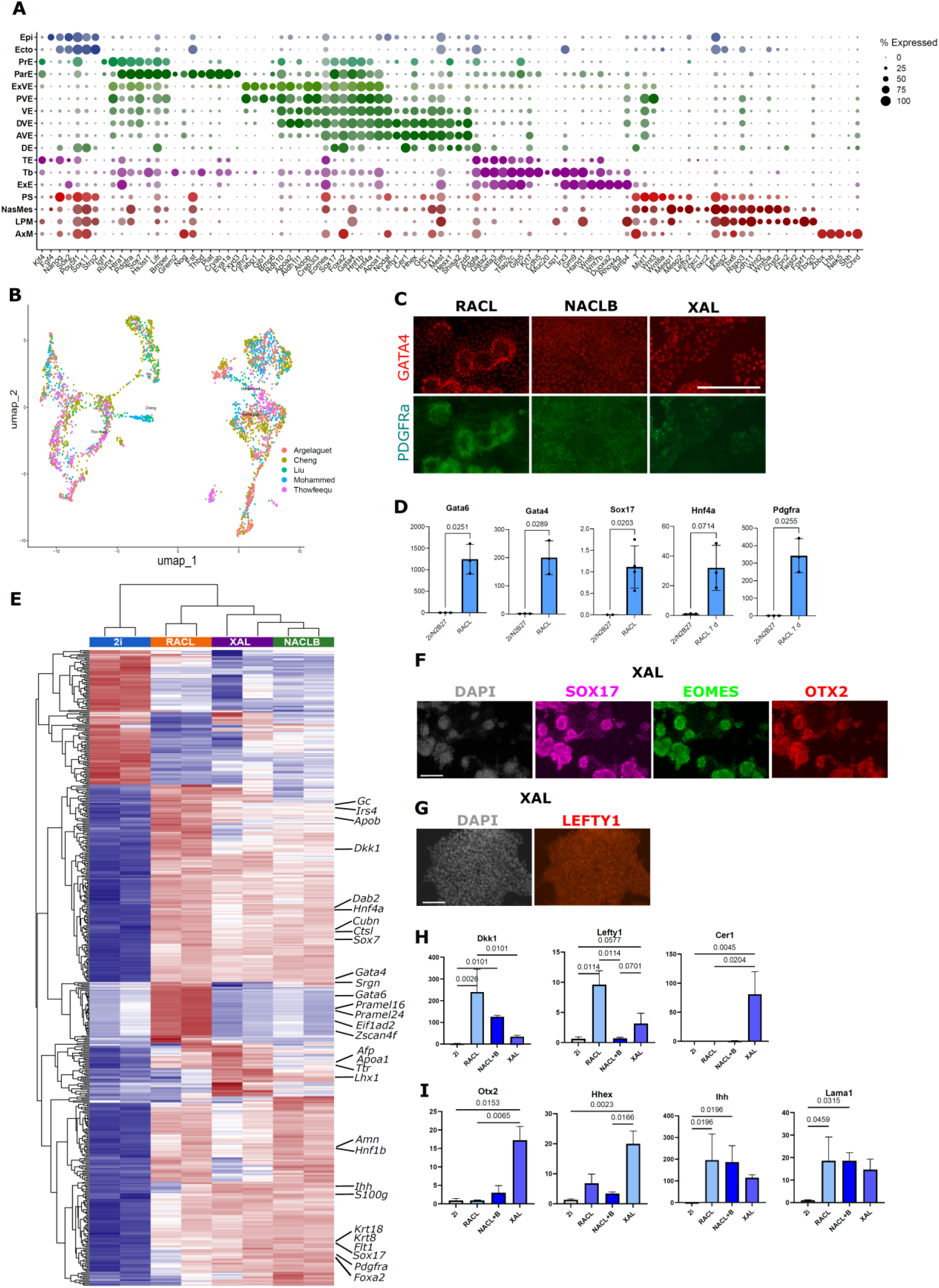
(A) Dot plot demonstrating lineage markers from differential expression analysis of integrated published datasets(Argelaguet et al., 2019; Cheng et al., 2019; Liu et al., 2022; Mohammed et al., 2017; Thowfeequ et al., 2024). (B) UMAP of integrated published mouse embryo development scRNAseq atlases (Argelaguet et al., 2019; Cheng et al., 2019; Liu et al., 2022; Mohammed et al., 2017; Thowfeequ et al., 2024) used as a reference. (C) Representative immunofluorescent images of RACL, NACLB and XAL 2D cultures. Scale bar: 100 µm. (D) Quantitative RT-PCR of gene expression in 2i/N2B27 and RACL cells. t-test with Welch’s correction (n=3, N=3). (E) Correlation based hierarchical clustering of top 500 genes expressed in 2i, RACL, NACLB and XAL cells based on bulk RNA-seq (n=2). (F-G) Representative immunofluorescent images of XAL 2D culture. Scale bar: 100 µm. (H-I) Quantitative RT-PCR expression analysis of 2i/N2B27, RACL, NACLB and XAL cells, non-parametrical Kruskal-Wallis test (n=3-5, N=3).

**Supplementary figure 2.**
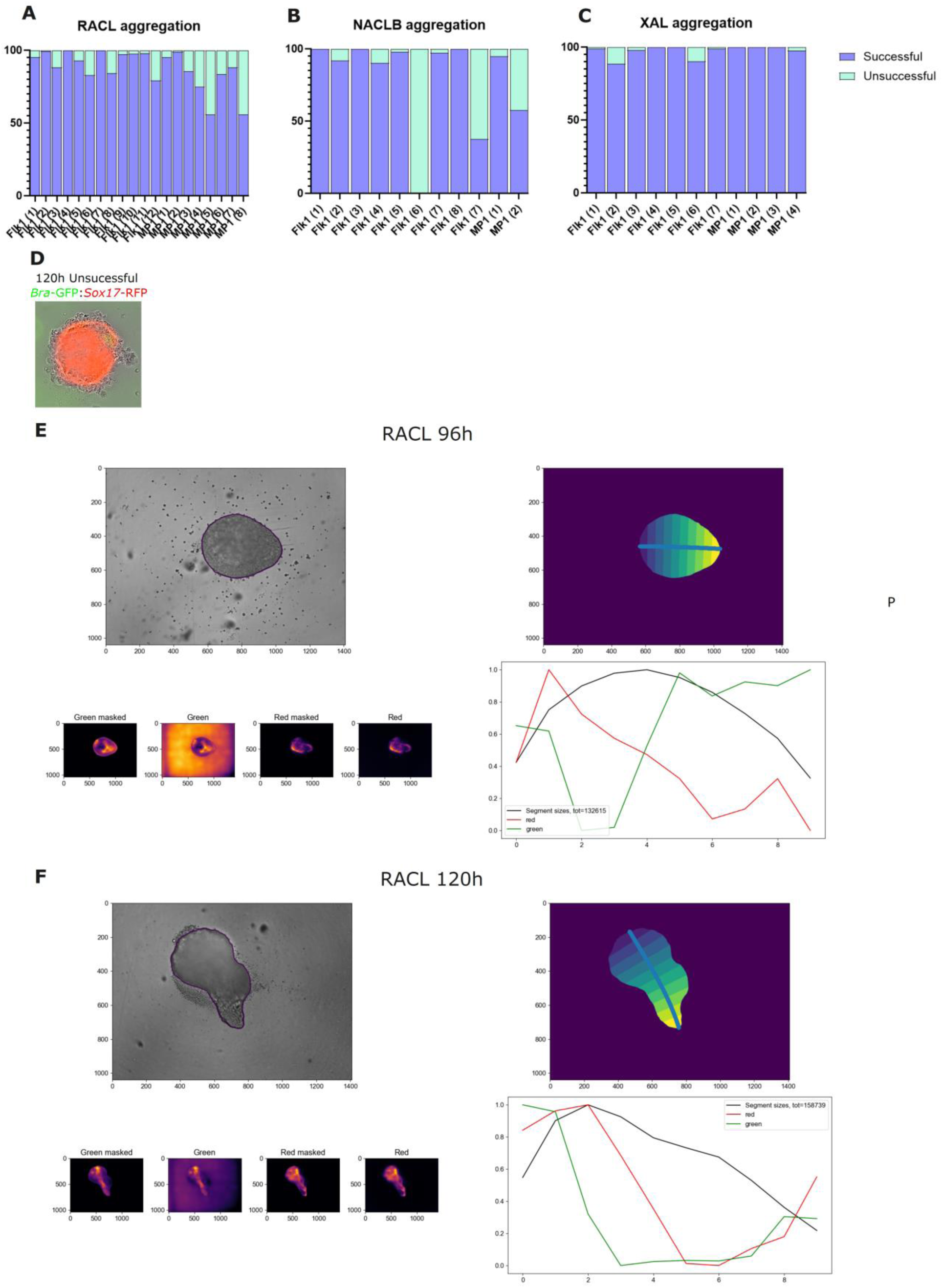
(A-C) Stacked plots showing the ratio of “successful” (elongated, symmetry break) and “non-successful” structures fully enveloped in ExEnd-like cells and failing to elongate in RACL (n=1859, 19 experimental replicates) (A), NACLB (n=616, 10 experimental replicates) (B), and XAL(n=923, 10 experimental replicates) (C) co-aggregation experiments. (D) A representative fluorescent image with *Sox17*-RFP and *Bra*-GFP overlay showing an example of “unsuccessful structure” at 120h. RACL cells don’t have fluorescent reporter, thus the signal is of gastruloid origin. (E-F) An example of morphometric and fluorescent signal measurements and masking used in Figure 2I-J (Methods) applied to 96h (E) and 120h (F) RACL-aggregoid.

**Supplementary figure 3.**
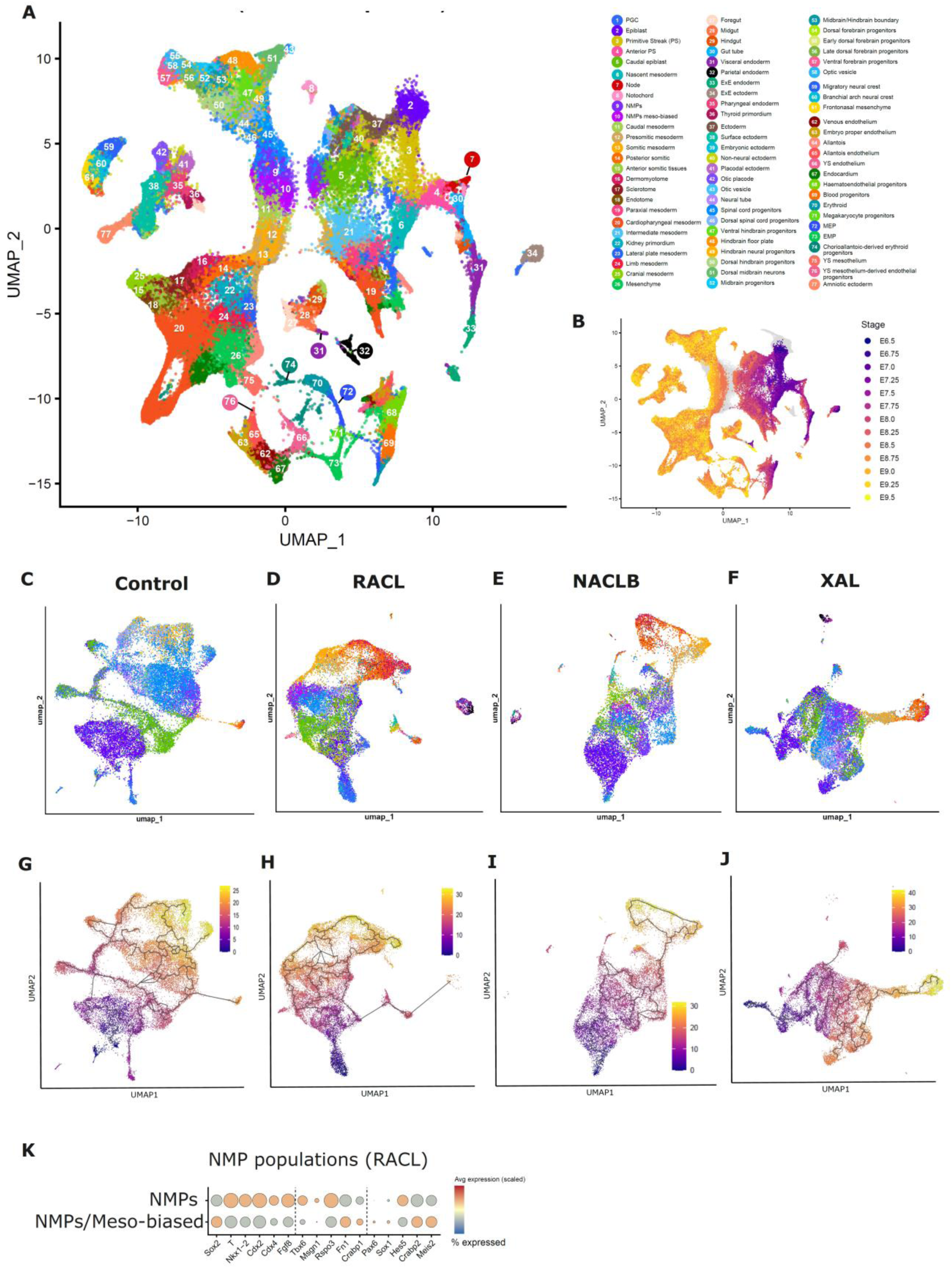
(A-B) UMAPs of scRNAseq datasets from the published mouse embryo E6.5-9.5 reference dataset(Imaz-Rosshandler et al., 2024; Pijuan-Sala et al., 2019). Color- and number-coded according to the common legend. (C-F) Integrated UMAP visualization of control gastruloids (C) and RACL(D), NACLB (E) and XAL(F)-aggregoids across time-points 48-120h. Color-coded according to the common legend. (G-J) Integrated UMAP visualization of control gastruloids (G) and RACL (H), NACLB (I) and XAL(J)-aggregoids across time-points 48-120h with pseudotime and trajectory projection (Monocle 3). (K) Dot-plot, z-scored, showing the relative differences of NMP, mesodermal and neuroectodermal markers expression in NMPs and NMPs mesoderm-biased clusters.

**Supplementary figure 4.**
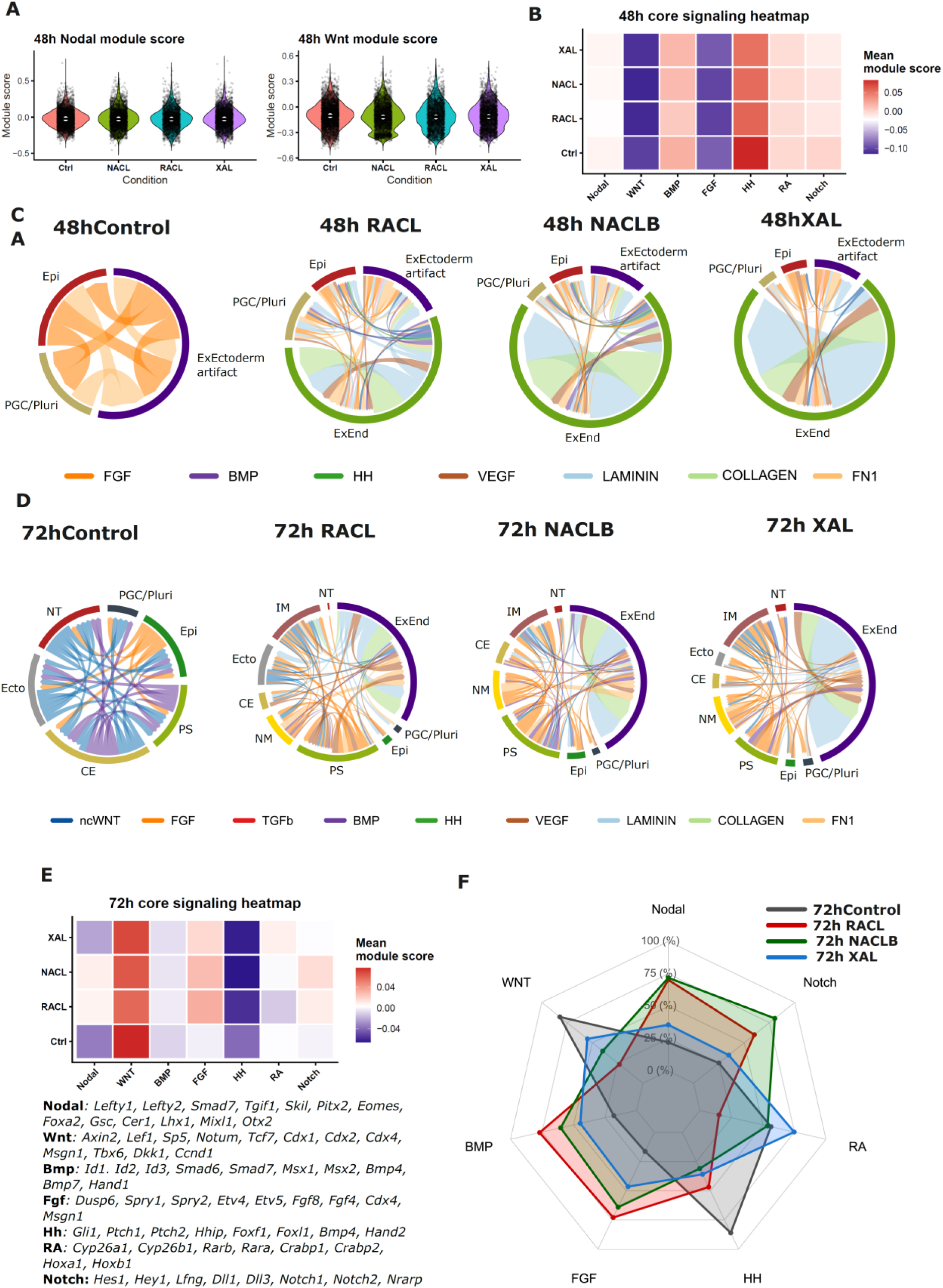
(A) Violin plots of NODAL and WNT scores (calculated expression of genes listed in Figure 4J) at 48h. (B, E) Heatmaps showing module scores of most selected signaling pathways: NODAL, WNT, BMP, NOTCH, RA, HH, FGF (list of genes is below) – at 48h (B) or at 72h (E) in all conditions. (C-D) Chord plots based on CellChat showing major communication pairs at 48h (C) or 72h (D). (F) Radar plot comparing z-scored module scores for NODAL, WNT, BMP, NOTCH, RA, HH, FGF at 72h.

**Movie 01.**

Time-lapse (1h resolution, 4 frames/second) video of control gastruloid development between 72 and 120h. Phase contrast, *Bra*-GFP: Sox17-RFP reporter line. Scale bar: 400 µm.

**Movie 02.**

Time-lapse (2h resolution, 4 frames/second) video of RACL-aggregoid development between 72 and 120h. Phase contrast, *Bra*-GFP: Sox17-RFP reporter line used for gastruloid part. Scale bar: 400 µm.

**Movie 03.**

Confocal time-lapse (15 minutes resolution) video of RACL-aggregoid development between 72 and 96h. *Bra*-GFP: Sox17-RFP reporter line used for gastruloid part. *Rosa26-*BFP*: Flk1-*GFP reporter line was used for RACL cells. BFP-positive cells represent the added ExEnd, RFP-positive cells correspond to DE produced by gastruloid. Scale bar: 100 µm.

**Movie 04.**

Brightfield video of RACL-aggregoid at 144h with contracting region corresponding to cardiac cells. Scale bar: 275 µm.

## Notes

### Competing Interest Statement

The authors have declared no competing interest.

### Summary of Updates

The current version of manuscript contains massive updates in quality of figures, scRNAseq analysis, image analysis, and cell communication analysis. The text was rewritten, including results and discussion. Funding information is also updated.

